# Small Molecule Activation of NAPE-PLD Enhances Efferocytosis by Macrophages

**DOI:** 10.1101/2023.01.25.525554

**Authors:** Jonah E. Zarrow, Abdul-Musawwir Alli-Oluwafuyi, Cristina M. Youwakim, Kwangho Kim, Andrew N. Jenkins, Isabelle C. Suero, Margaret R. Jones, Zahra Mashhadi, Kenneth P. Mackie, Alex G. Waterson, Amanda C. Doran, Gary A. Sulikowski, Sean S. Davies

## Abstract

*N*-acyl-phosphatidylethanolamine hydrolyzing phospholipase D (NAPE-PLD) is a zinc metallohydrolase that hydrolyzes *N*-acyl-phosphatidylethanolamine (NAPEs) to form *N*-acyl-ethanolamides (NAEs) and phosphatidic acid. Several lines of evidence suggest that reduced NAPE-PLD activity could contribute to cardiometabolic diseases. For instance, *NAPEPLD* expression is reduced in human coronary arteries with unstable atherosclerotic lesions, defective efferocytosis is implicated in the enlargement of necrotic cores of these lesions, and NAPE-PLD products such as palmitoylethanolamide and oleoylethanolamide have been shown to enhance efferocytosis. Thus, enzyme activation mediated by a small molecule may serve as a therapeutic treatment for cardiometabolic diseases. As a proof-of-concept study, we sought to identify small molecule activators of NAPE-PLD. High-throughput screening followed by hit validation and primary lead optimization studies identified a series of benzothiazole phenylsulfonyl-piperidine carboxamides that variably increased activity of both mouse and human NAPE-PLD. From this set of small molecules, two NAPE-PLD activators (VU534 and VU533) were shown to increase efferocytosis by bone-marrow derived macrophages isolated from wild-type mice, while efferocytosis was significantly reduced in *Napepld*^*-/-*^ BMDM or after Nape-pld inhibition. Together these studies demonstrate an essential role for NAPE-PLD in the regulation of efferocytosis and the potential value of NAPE-PLD activators as a strategy to treat cardiometabolic diseases.

## Introduction

Atherosclerotic Cardiovascular Disease (ASCVD) remains a major cause of death in the United States and throughout the world. Efferocytosis, the non-inflammatory clearance of apoptotic cells by macrophages, is a critical step in the resolution of inflammation and defective efferocytosis has been implicated in the development and expansion of necrotic cores within atherosclerotic plaques^1-2^. Plaques with large necrotic cores are highly vulnerable to rupture, thereby triggering potentially fatal thrombosis and myocardial infarction^3^. Therefore, identifying the factors whose dysfunction impairs efferocytosis and appropriate countermeasures against this dysfunction could lead to novel treatments for ASCVD.

NAPE-PLD is a zinc metallohydrolase within the metallo-β-lactamase superfamily^4-5^. NAPE-PLD hydrolyzes *N*-acyl-phosphatidylethanolamines (NAPEs) to phosphatidic acid and *N-*acyl-ethanolamides (NAEs) such as palmitoylethanolamide (PEA) and oleoylethanolamide (OEA)^4-5^ **(Figure 1)**. In rodents, high-fat or Western diets decrease *Napepld* expression and levels of PEA and OEA in a variety of tissues^6-8^. *NAPEPLD* expression is also reduced in atherosclerotic plaques (especially unstable plaques) of human coronary arteries ^8^. In mice, directly administering NAEs or NAE-boosting bacteria counteract atherosclerosis as well as other cardiometabolic diseases including obesity, glucose intolerance, and non-alcoholic fatty liver disease^8-12^. Importantly, these treatments inhibit enlargement of the necrotic core within atherosclerotic lesions^8, 10-11^. Cellular studies show that PEA and OEA enhance M2 polarization and the efferocytosis capacity of bone-marrow derived macrophages (BMDM) via GPR55 and/or PPARα dependent mechanisms^8 10^. Together these studies suggest that reduced macrophage *NAPEPLD* expression could lead to reduced efferocytosis by macrophages and thereby drive expansion of the necrotic core and atherosclerosis.

**Figure 1.**
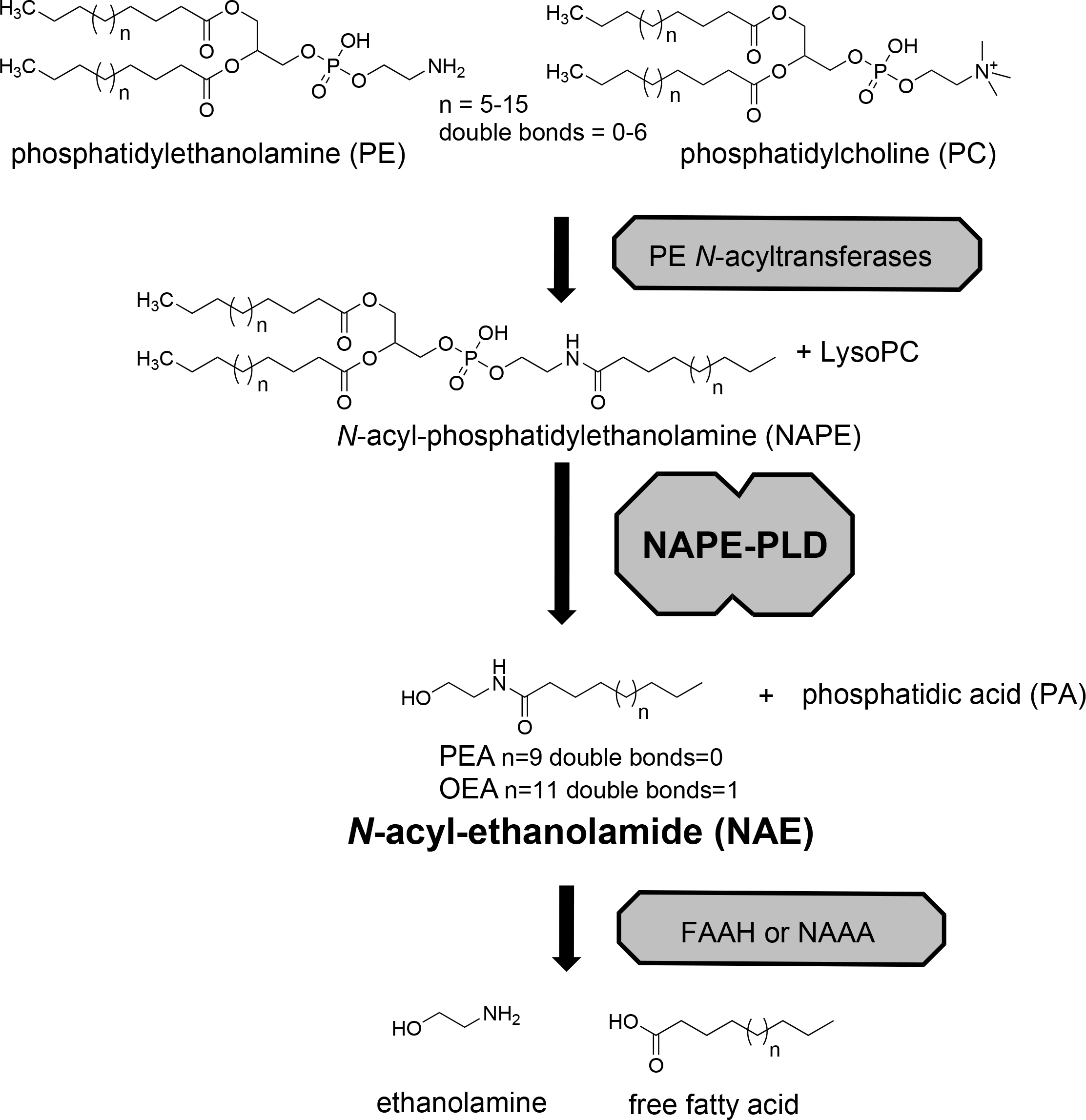
Biosynthesis of *N*-acyl-ethanolamides via NAPE-PLD. *N*-acyl-ethanolamides (NAEs) including palmitoylethanolamide (PEA) and oleoylethanolamide (OEA) are formed by NAPE-PLD pathway in a two step process. First, PE N-acyltransferases transfer an acyl chain from phosphatidylethanolamine (PC) to the nitrogen of phosphatidylethanolamine (PE) to generate N-acyl-phosphatidylethanolamines (NAPEs), and lysophosphatidylcholine (lysoPC). Then NAPE-PLD cleaves NAPE at the distal phosphodiester bond to generate the NAE and phosphatidic acid (PA). NAEs then act on receptors including PPARa, GPR119, and GPR55 to exert biological effects. NAEs are rapidly inactivated by fatty acid amide hydrolase (FAAH) and *N*-acylethanolamide acid amidase (NAAA) by their degradation to ethanolamine and free fatty acid.

If NAPE-PLD regulates efferocytosis, then small molecules that enhance macrophage enzyme activity should enhance macrophage efferocytosis and could therefore potentially inhibit the development of unstable atherosclerotic lesions. While several small molecule inhibitors of NAPE-PLD have been reported recently^13-15^, there are currently no small molecule activators of NAPE-PLD. We therefore sought to identify small molecules that could enhance macrophage enzyme activity in order to test their effects on the macrophage efferocytosis capacity.

## Results

### New NAPE-PLD activator chemotype identified by HTS and early SAR studies

We screened 39,328 compounds from the Vanderbilt Discovery Collection, a chemical library of lead-like compounds, for their effects on Nape-pld activity using the commercially available fluorogenic NAPE analog, PED-A1 (**Figure 2A**) and recombinant mouse Nape-pld^14^. The change in the measured rate of fluorescence after addition of commercially available PED-A1 (**Figure 2B**) was used to calculate a B-score for each compound (**Figure 2C**). From the 39,238 compounds screened, 399 were judged as potential activators based on observed change in fluorescence) by at least 3 standard deviations compared to vehicle.

**Figure 2.**
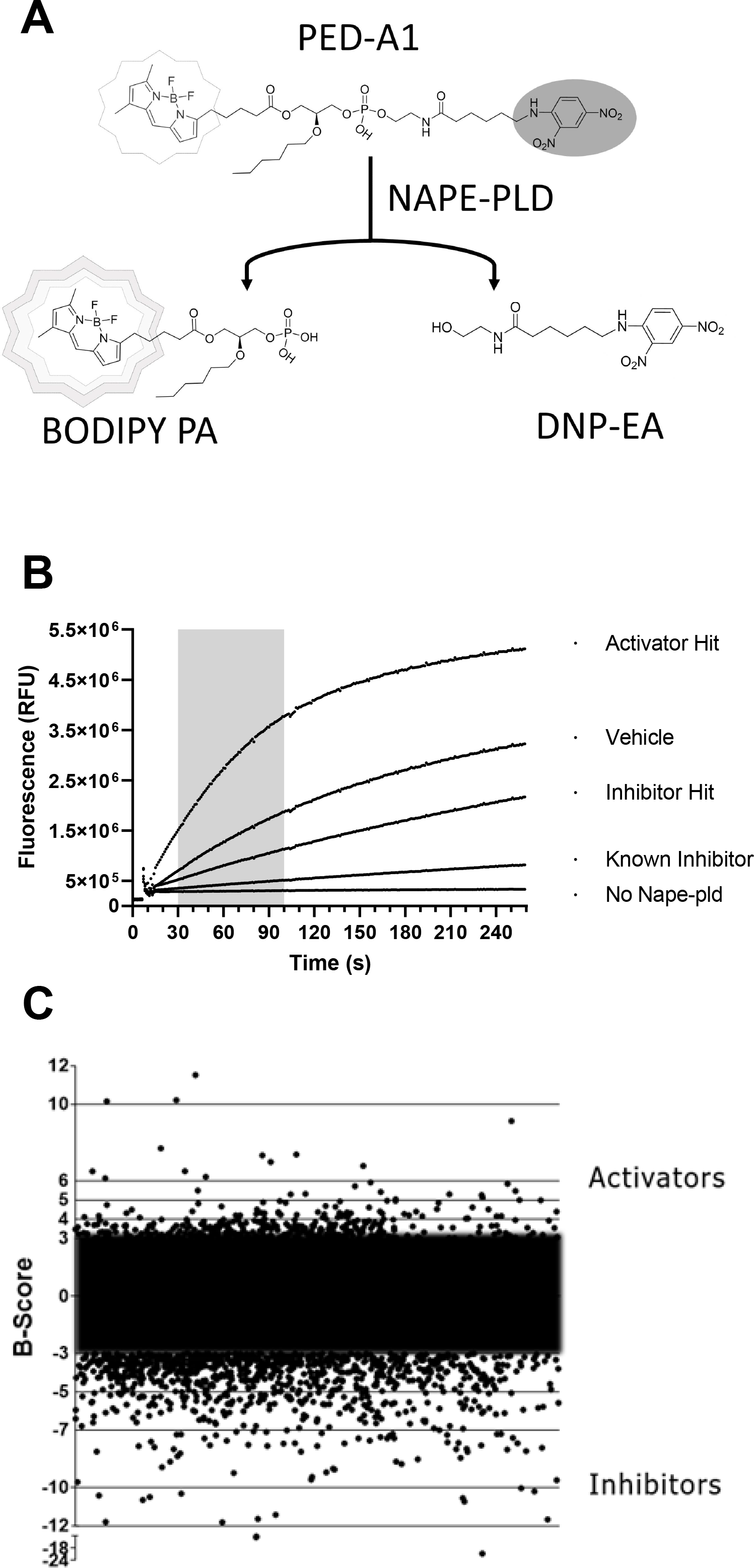
High throughput screening identifies potential NAPE-PLD activators. **A.** Schematic representation of the HTS assay which detects Nape-Pld activity by the fluorescence resulting from the hydrolysis of the fluorogenic NAPE analog PED-A1. Intact PED-A1 only weakly fluoresces due to internal quenching by its dinitrophenyl moiety. Nape-pld hydrolysis of PED-A1 generates dinitrophenyl-hexanoyl-ethanolamide (DNP-EA and highly fluorescent BODIPY-labeled phosphatidic acid (BODIPY-PA). **B**. Sample activity curves from HTS for controls, a representative activator hit and a representative inhibitor hit. The shaded region represents the time period used in scoring fluorescence changes for all compounds. **C**. B-scores of various test library compounds in the HTS assay. Compounds with B-scores ≥ 3 (activators) or ≤ -3 (inhibitors) were deemed to be potential hits.

Three of the 399 identified activators (**Table 1**, entries 1-3) shared a common benzothiazole phenylsulfonyl-piperidine carboxamide (BT-PSP) core structure. To determine if this series of benzothiazoles might serve as a tractable lead for the development of activators as chemical probes, we obtained additional structurally similar compounds using a combination of targeted purchasing of commercial molecules (**Table 1**, entries 4-13) and discrete chemical synthesis (**Table 1**, entries 14-22).

**Table 1.** Preliminary SAR of NAPE-PLD activators^a^. **In vitro Nape-pld modulation by benzothiazole phenylsulfonyl-piperidine carboxamides (BT-PSPs)**. EC_50_ represent concentration (in μM) required for half-maximal activity, expressed in μM. E_max_ represents maximal increase in activity (as fold activity of vehicle control).

| Entry | Structure | VU Number | EC<sub>50</sub> μM<sup>b</sup><br>(95% CI) | E<sub>max</sub><sup>c</sup><br>(95% CI) |
| --- | --- | --- | --- | --- |
| 1 |  | VU484 | 2.3<br>(1.8-3.1) | 2.1<br>(2.0-2.2) |
| 2 |  | VU485 | 12.0<br>(7.3-85.9) | 2.0<br>(1.8-4.0) |
| 3 |  | VU486 | 0.32<br>(0.15-0.58) | 1.8<br>(1.7-1.9) |
| 4 |  | VU0488 | 3.3<br>(2.7-4.2) | 1.7<br>(1.6-1.8) |
| 5 |  | VU539 | 0.74<br>(0.46-1.1) | 1.7<br>(1.6-1.8) |
| 6 |  | VU542 | 0.95<br>(0.66-1.36) | 1.9<br>(1.9-2.0) |
| 7 |  | VU517 | 1.1<br>(0.9-1.4) | 2.0<br>(2.0-2.1) |
| 8 |  | VU534 | 0.30<br>(0.17-0.47) | 2.1<br>(2.0-2.2) |
| 9 |  | VU533 | 0.30<br>(0.16-0.44) | 2.6<br>(2.6-2.8) |
| 10 |  | VU536 | 1.1<br>(0.7-1.6) | 2.3<br>(2.2-2.4) |
| 11 |  | VU575 | 0.57<br>(0.34-0.92) | 2.4<br>(2.3-2.5) |
| 12 |  | VU0601 | 0.94<br>(0.71-1.2) | 1.8<br>(1.8-1.9) |
| 13 |  | VU605 | 0.74<br>(0.50-1.22) | 2.1<br>(2.0-2.2) |
| 14 |  | VU231 | 1.1<sup>d</sup> | 2.0<sup>d</sup> |
| 15 |  | VU205 | NA<sup>e</sup> | <1.2 |
| 16 |  | VU212 | 2.5<br>(1.8-3.3) | 1.6<br>(1.5-1.6) |
| 17 |  | VU233 | NA<sup>e</sup> | <1.2 |
| 18 |  | VU209 | 3.5<br>(2.4-5.4) | 1.9<br>(1.8-2.1) |
| 19 |  | VU227 | 4.1<br>(2.6-9.1) | 1.7<br>(1.6-1.9) |
| 20 |  | VU210 | 1.8<br>(1.3-2.3) | 2.4<br>(2.3-2.6) |
| 21 |  | VU203 | 4.8<br>(3.2-7.4) | 1.8<br>(1.7-2.0) |
| 22 |  | VU211 | 1.4<br>(1.0-1.9) | 2.1<br>(2.0-2.2) |

<sup>a</sup>Compounds were either purchased from commercial source or prepared in house by chemical synthesis. <sup>b</sup>EC<sub>50</sub>: concentration needed for 50% maximal activator effect. <sup>c</sup>E<sub>max</sub>: maximal efficacy expressed as fold increase in Nape-pld activity versus vehicle only. <sup>d</sup>95% CI for EC<sub>50</sub> and E<sub>max</sub> indeterminant due to poor fit. <sup>e</sup>NA: not active.

The activity of this series of 22 benzothiazole phenylsulfonyl-piperidine carboxamides thus obtained was assessed for enzyme activation using recombinant mouse Nape-pld (**Supplemental Figure 1 and Table 1**). Compounds **VU534** and **VU533** (entries 8 and 9) proved the most potent of the series of enzyme activators, both showing half-maximal activation concentrations (EC_50_) of 0.30 μM and a more than two-fold maximal induction of Nape-pld activity relative to vehicle controls (E_max_ > 2.0). Nine other BT-PSPs (entries 3, 5-7, 10-14) had EC_50_ ≤ 1.1 μM and E_max_ > 1.7. The remaining members of the series had EC_50_ < 10 μM and E_max_ > 1.7, except that compound **VU212** (entry 16) showed poor efficacy (E_max_ < 1.6), and **VU205** and **VU233** proved inactive (Emax <1.2). With this preliminary structure-activity relationship (SAR) data, we selected **VU534** and **VU533** as chemical probes for study of NAPE-PLD activation and **VU233** as a negative control in studies outlined below.

### Utility as probes of NAPE-PLD activation in mouse and human cells

To determine if the selected small molecules could be used as probes of NAPE-PLD activation in cultured cells, we first evaluated their cytotoxicity in RAW264.7 mouse macrophages and HepG2 human hepatocytoma cells. Graded concentrations of **VU534** showed minimal cytotoxicity up to 30 μM in either cell line **(Supplemental Figure 2A** and **2B**). Likewise, **VU533** and **VU233** also showed no cytotoxicity when tested at 30 μM in either RAW264.7 (**Supplemental Figure 2C**) or HepG2 cells **(Supplemental Figure 2D**).

We next examined the efficacy of activators **VU534** and **VU533** and inactive **VU233** to increase Nape-pld activity in RAW264.7 cells. Both activators **VU534** and **VU533** significantly increased Nape-pld activity, while **VU233** showed no significant effect (**Figure 3A**). Other analogs in the series of 22 benzothiazole phenylsulfonyl-piperidine carboxamides that increased the activity of recombinant Nape-pld in the biochemical assay (**Table 1**) also significantly increased Nape-pld activity in RAW264.7 cells in a concentration-dependent manner (**Supplementa**l **Figure 3**), with a correlation observed between the efficacy in the biochemical assay and their efficacy in RAW264.7 cells (**Figure 3B**). To confirm that the effect seen in RAW264.7 cells was due to Nape-pld modulation, we tested the effects of bithionol, a known irreversible inhibitor of NAPE-PLD^14^. The increase in RAW264.7 cellular Nape-pld activity induced by 20 μM **VU534** was blocked in a concentration-dependent manner by bithionol (**Figure 3C**). These results are consistent with enzyme activation of **VU534** on PED-A1 hydrolysis being dependent on Nape-pld rather than an alternate modulating pathway.

**Figure 3.**
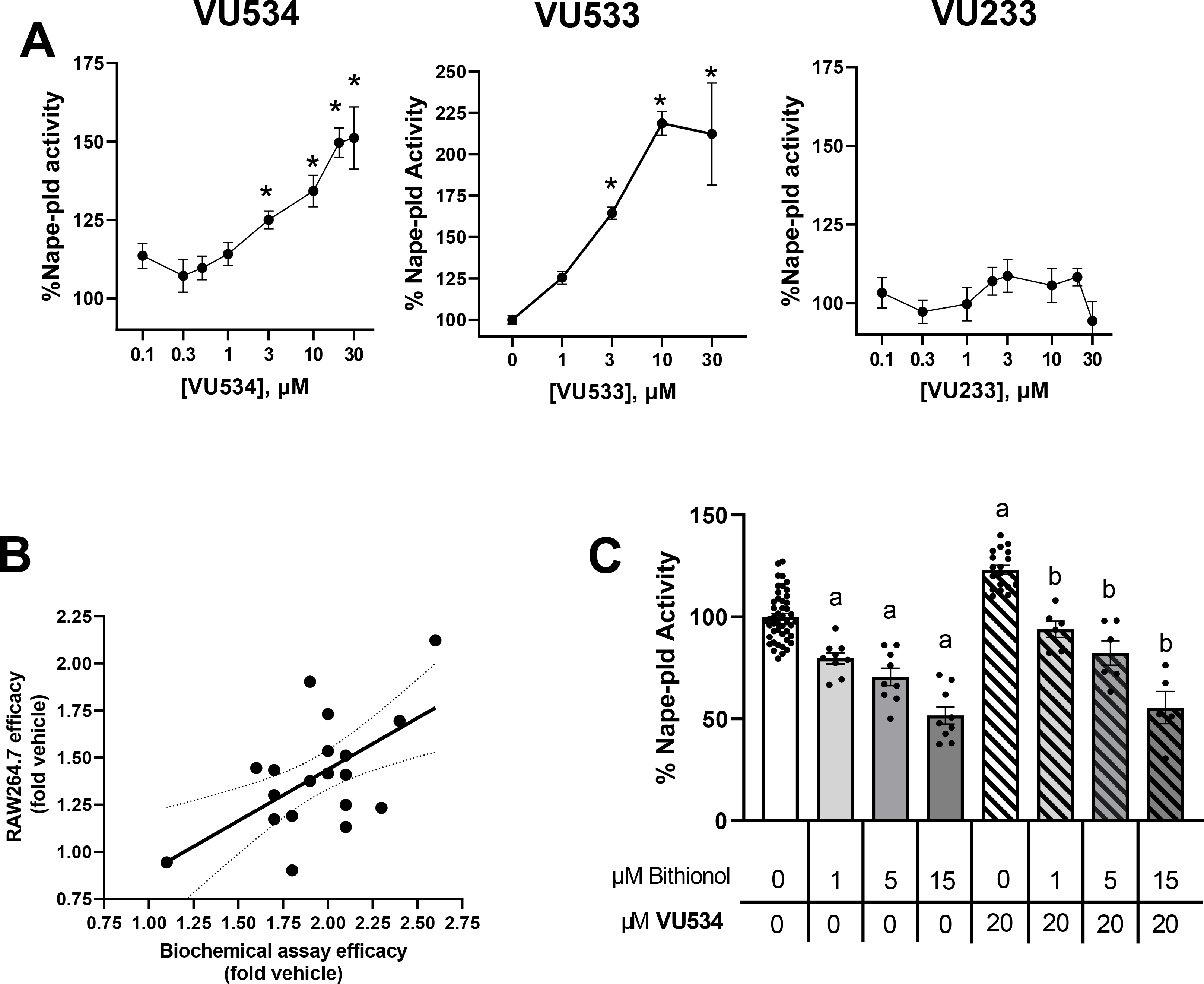
VU534 and VU533 increase the NAPE-PLD activity of RAW264.7 macrophages. **A.** Effect of graded concentrations of compound **8** (left panel), compound **9** (middle panel), and compound **17** (right panel) on Nape-pld activity in RAW264.7 cells, measured using PED-A1. Each compound was tested on at least two separate days and the individual replicates from each day normalized to vehicle control and then combined (mean ± SEM, n = 4-11). 1-way ANOVA p<0.0001 for **VU534**, p<0.0001 for **VU533**, and p=not significant for **VU233**. *p<0.05 vs 0 μM, Dunnet’s multiple comparison test. **B**. Correlation between maximal efficacy of 19 BT-PSPs and analogs in the recombinant Nape-pld assay and their efficacy at 30 μM in cultured RAW264.7 cells. Simple linear regression. Slope=0.5455, R^2^ 0.3394 p=0.0089 for slope significantly non-zero. **C**. Bithionol, a Nape-pld inhibitor, blocks increased Nape-pld activity in RAW264.7 cells induced by compound **8**. 1-way ANOVA p<0.0001 Sidak’s multiple comparison test, ^a^ p<0.05 vs 0 μM Bith with 0 μM **VU534** group, ^b^p<0.05 vs 0 μM Bith with 20 μM **VU534** group.

We then determined the effects of NAPE-PLD modulators on purified human NAPE-PLD and human-derived culture cells. Both **VU534** and **VU533** invoked concentration-dependent increases in the activity of recombinant human NAPE-PLD, while the inactive **VU233** had no significant effect (**Figure 4A**). The potency of **VU534** and **VU533** for activating human NAPE-PLD was somewhat less than for activating mouse Nape-pld **(Table 1**). Other benzothiazoles (**Table 1**) also activated recombinant human NAPE-PLD (**Supplemental Figure 4**). **VU534** and **VU533**, but not the inactive compound **VU233**, increased NAPE-PLD activity in HepG2 cells in a concentration-dependent manner (**Figure 4B**).

**Figure 4.**
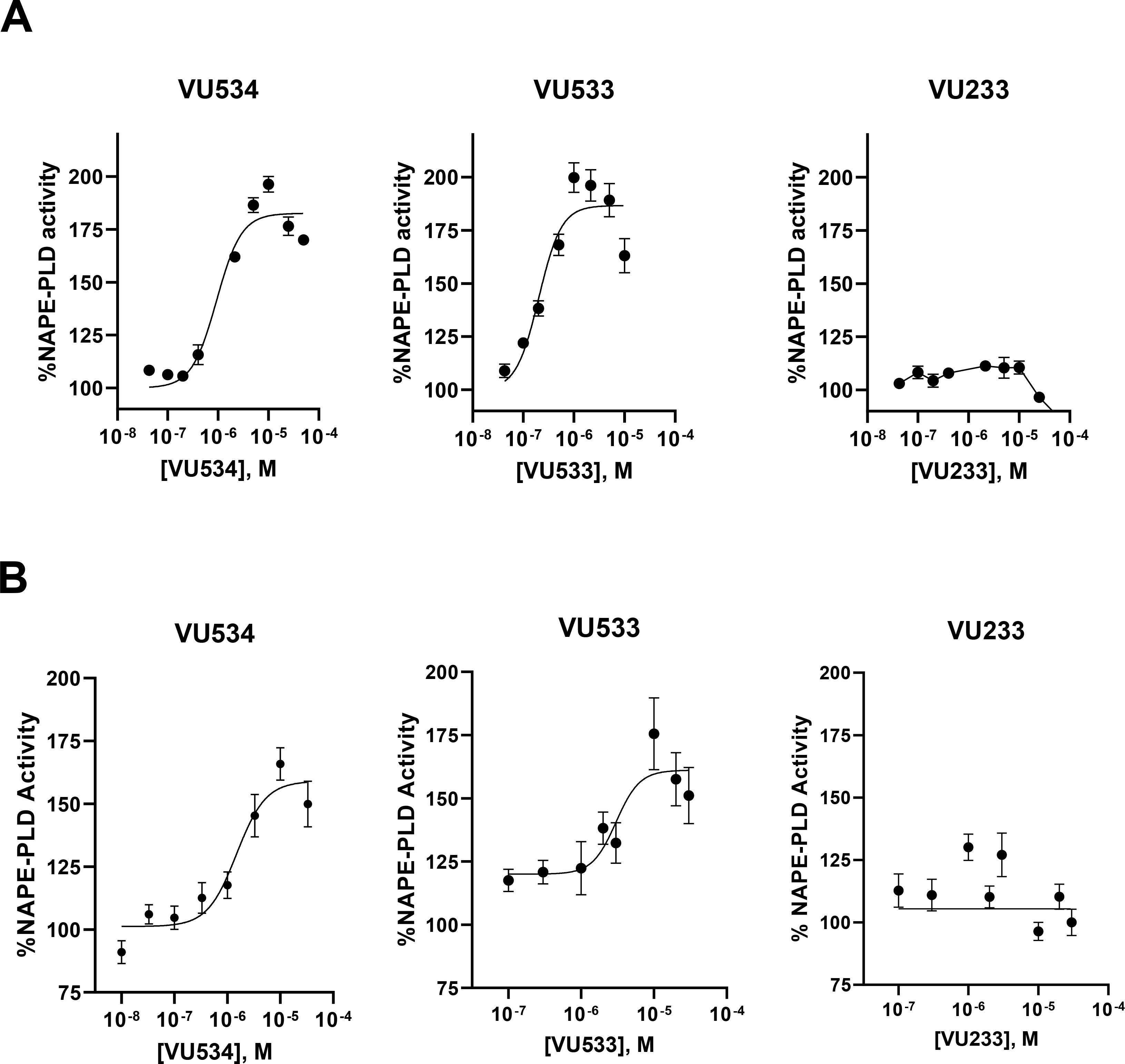
VU534 and VU533 activate human NAPE-PLD. **A.** Effect of graded concentrations of **VU534, VU533**, and **VU233** on activity of recombinant human NAPE-PLD with PED-A1 as substrate. Mean ± SEM, n = 3. Non-linear regression with variable slope (four parameter) was used to calculate EC_50_ and E_max_. **VU534** EC_50_ 0.93 μM (95% CI 0.63 to 1.39 μM), E_max_ 1.8-fold (95% CI 1.8 to 1.9-fold); **VU533** EC_50_ 0.20 μM (95% CI 0.12 to 0.32 μM),E_max_ 1.9-fold (95% CI 1.8 to 2.0-fold); **VU233** not calculable. **B**. Effect of graded concentrations of **VU534, VU533**, and **VU233** on NAPE-PLD activity of HepG2 cells measured using flame-NAPE as substrate. Each compound was tested on two separate days and the individual replicates from each day normalized to vehicle control and then combined (mean ± SEM, n = 4-6); **VU534** EC_50_ 1.5 μM (95% CI 0.6 to 2.8 μM), E_max_ 1.6-fold activity (95% CI 1.5 to 1.8-fold); **VU533** EC_50_ 3.0 μM (1.4 to 5.7 μM), E_max_ 1.6-fold (95% CI 1.5 to1.8) fold; **VU233** not calculable.

### Biochemical characterization of lead NAPE-PLD activators

To confirm the results of our fluorescence-based assays, we employed an orthogonal biochemical Nape-pld assay based on LC/MS, similar to those employed previously^13, 16-17^. Recombinant mouse Nape-pld was pre-treated with **VU534, VU533, VU233** or vehicle for 30 min, then *N*-oleoyl-phosphatidylethanolamine (NOPE) was added for 90 min, and the resulting levels of OEA and NOPE was measured by LC/MS. Both activators **VU534** and **VU533** significantly increased the OEA/NOPE ratio, while inactive **VU233** had no significant effect (**Figure 5A**). To further characterize the effect of **VU534** on Nape-pld activity, we performed a Michaelis-Menten analysis using recombinant mouse Nape-pld and graded concentrations of flame-NAPE, a selective NAPE-PLD fluorogenic substrate resistant to competing esterase activity^18^. Compound **VU534** lowered the K_1/2_ (12.4 μM with vehicle vs 5.9 μM with **VU534**) and increased maximal velocity (227 RFU/min with vehicle vs 421 RFU/min with **VU534**) of Nape-pld (**Figure 5B**).

**Figure 5.**
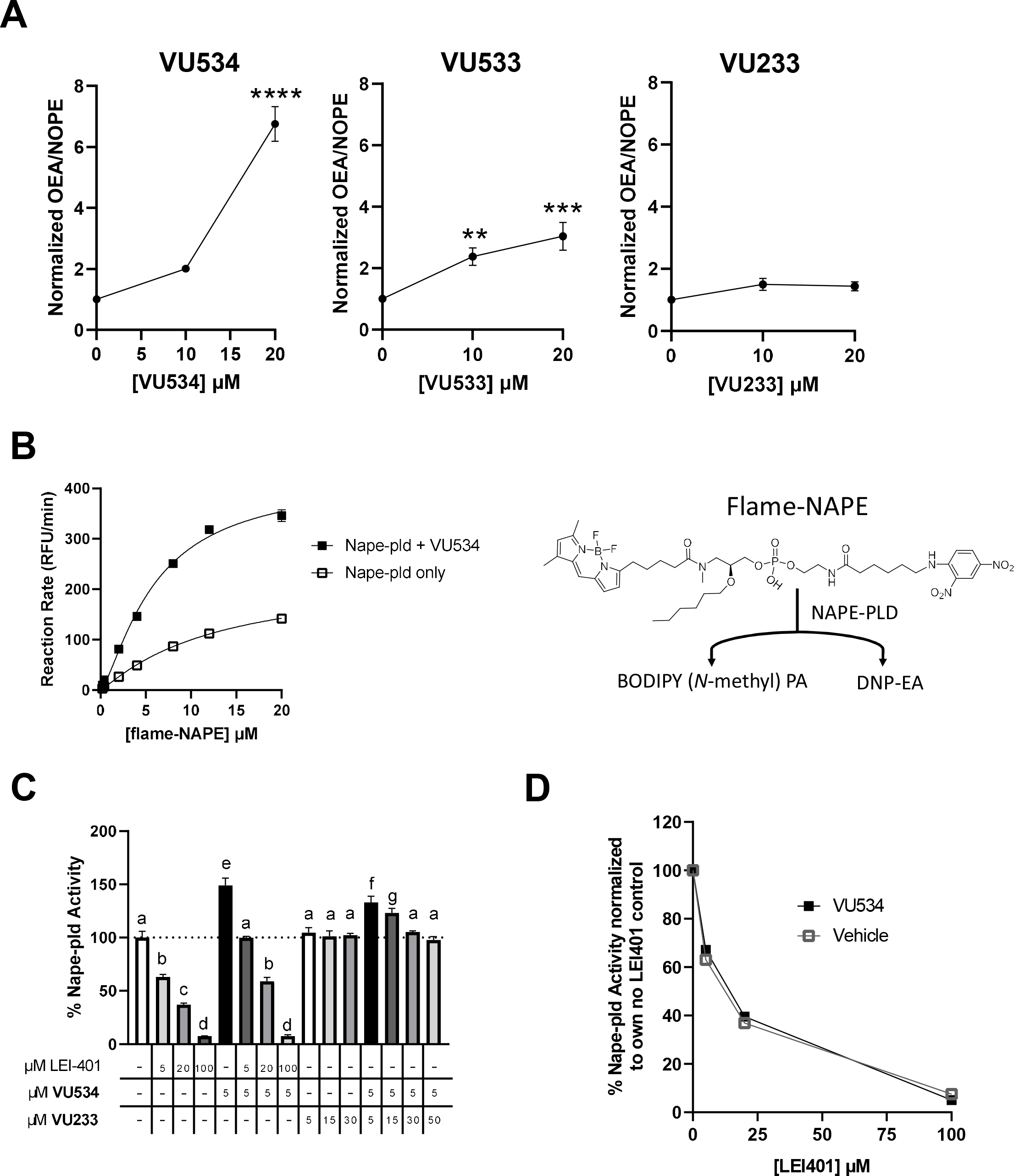
Additional characterization of Nape-pld modulation by VU534. **A.** Activity of recombinant mouse Nape-pld using *N*-oleoyl-phosphatidylethanolamine (NOPE) as substrate and measuring OEA and NOPE by LC/MS/MS. Ratio of OEA to NAPE was normalized to 0 μM compound control. The assays of **VU534** and **VU233** were performed using the same 0 μM compound replicates. 1-way ANOVA **VU534** p<0.0001, **VU533** p=0.007, **VU233** p = 0.0547; Dunnett’s multiple comparison test for individual compounds **p=0.0074, ***p=0.0005, ****p<0.0001 **B**. Michaelis-Menten analysis using flame-NAPE as substrate for recombinant mouse Nape-pld with or without **VU534**. Non-linear regression curves (allosteric sigmoidal) were used to calculate K_1/2_ and Vmax. **C**. In vitro competition assay for effects on flame-NAPE hydrolysis by recombinant mouse Nape-pld. 1-way ANOVA, p<0.0001. Groups sharing letters do not significantly differ from each other in Tukey multiple comparisons test. **D**. Data from in vitro competition assay data normalized to the value for 0 μM LEI-401 with 5 μM **VU534** for all samples with 5 μM **VU534** (VU534) or 0 μM LEI-401 with vehicle for all samples with no **VU534** (Vehicle). Samples treated with the same concentration of LEI-401, with or without 5 μM **VU534**, did not significantly differ, Sidak’s multiple comparisons test.

These results were consistent with **VU534** leading to NAPE-PLD activation by way of allosteric modulation. We therefore examined the effect of LEI-401 (a known, reversible inhibitor of NAPE-PLD)^13^ or the inactive **VU233** on the ability of **VU534** to enhance Nape-pld activity. LEI-401 inhibited mouse NAPE-PLD activity in a concentration-dependent manner, with a concentration of 100 μM achieving nearly complete inhibition (**Figure 5C**). In the absence of LEI-401, 5 μM **VU534** increased Nape-pld activity 1.5-fold. When 5 μM of compound **VU534** was added, LEI-401 still inhibited Nape-pld activity in a concentration-dependent manner, with 100 μM of LEI-401 again being sufficient for near complete effect. Although the absolute Nape-pld activity was higher in the presence of **VU534** than without it, when activity was normalized as percentage of initial activity, the extent of inhibition by LEI-401 was not significantly different (**Figure 5D**), suggesting that **VU534** does not compete with LEI-401 for the same binding site. Further insight into the mode of enzyme activation was obtained when inactive **VU233** was observed to suppress the enzyme activation of 5 μM **VU534** in a concentration-dependent manner (**Figure 5C**). These results are consistent with inactive **VU233** being a neutral allosteric binder that binds to the same site as **VU534**.

A survey of the literature revealed that a series of benzothiazoles with structural features somewhat similar to **VU534** and **VU533** have been developed as dual inhibitors of fatty acid amide hydrolase (FAAH) and soluble epoxide hydrolase (sEH)^19^. Because inhibition of FAAH would increase cellular NAE levels independently of NAPE-PLD (**Figure 1**), we tested whether our activators modulated FAAH activity. Graded concentrations of **VU534, VU533**, and **VU233** showed only weak inhibition of FAAH activity (**Figure 6A**). We also tested whether our activators modulated sEH activity. While sEH does not lie in the biochemical pathway for NAE biosynthesis or metabolism, inhibition or genetic ablation of sEH increases the levels of epoxy fatty acids^20^ and thereby exerts biological effects similar to the known effects of NAEs such as reducing obesity, cardiovascular disease, pain, and inflammation^20-23^. Graded concentrations of **VU534** showed modest inhibition of sEH (IC_50_ 1.2 μM, 95% CI 0.5-2.4 μM, maximal inhibition 55%), while neither **VU533** nor **VU233** significantly inhibited seH **(Figure 6B)**. Other compounds of our series showed variable effects on FAAH and sEH activity (**Supplemental Figure 5** and **6**). These modest off-target effects should not significantly interfere with the use of these compounds to probe the contribution of Nape-pld activity as long as bona fide sEH inhibitors are also tested as controls.

**Figure 6.**
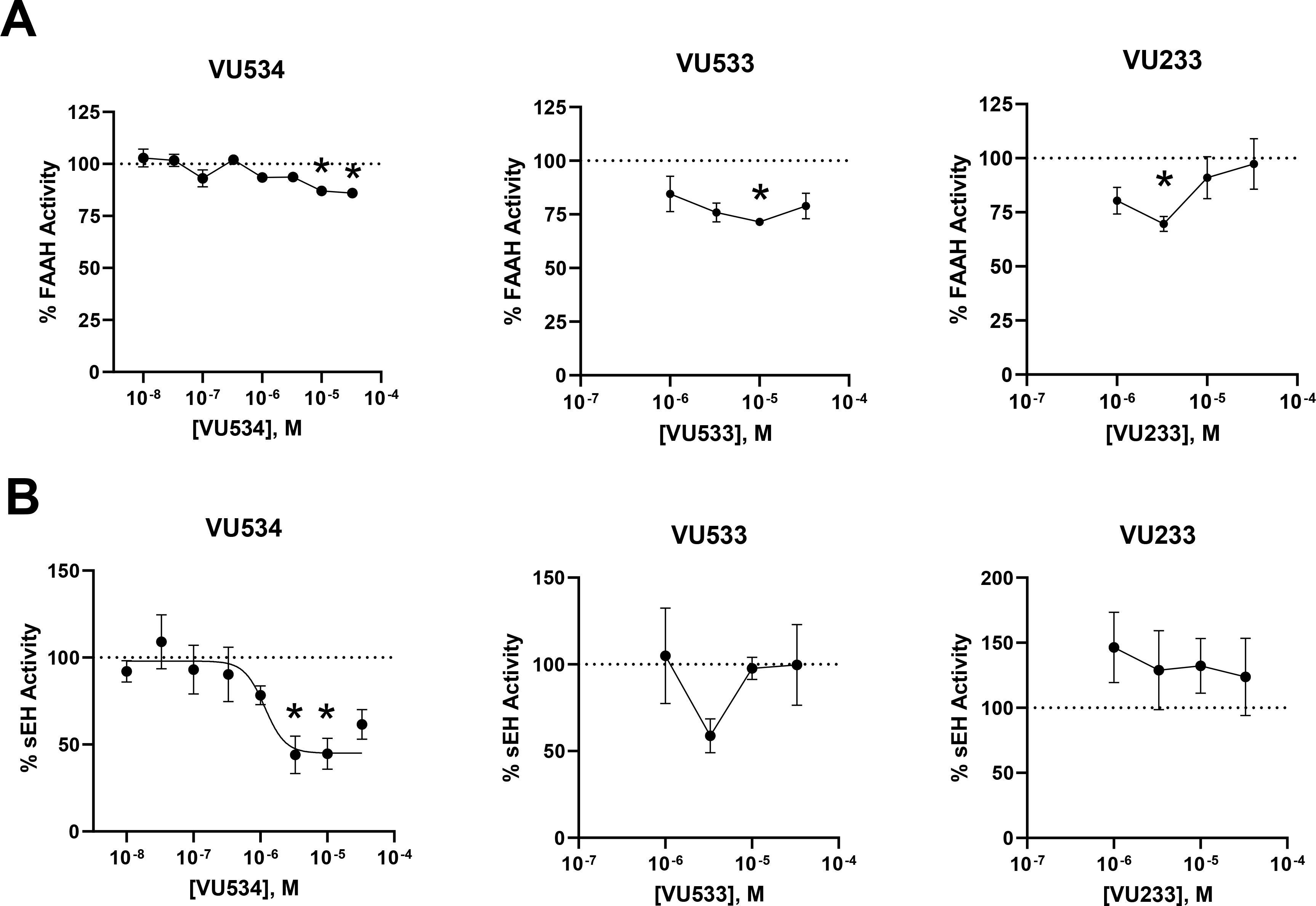
Evaluation of off-target effects on FAAH and sEH. **A.** Effects of graded concentrations of **VU534, VU533**, or **VU233** on activity of Fatty Acid Amide Hydrolase (FAAH). 1-way ANOVA VU534 p=0.0007, VU533 p<0.0001, and VU233 p <0.0001;*p<0.05 vs 0 μM, Dunnett’s multiple comparison test for individual compounds **B**. Effects of graded concentrations of **VU534, VU533**, or **VU233** on activity of soluble Epoxide Hydrolase (sEH). 1-way ANOVA **VU534** p<0.0001, **VU533** p<0.0001, and **VU233** p<0.0001; *p<0.05 vs 0 μM, Dunnett’s multiple comparison test for individual compounds.

### Modulation of Nape-pld activity modulates efferocytosis by macrophages

*Rinne et al* showed that PEA, a Nape-pld product, enhanced the ability of bone-marrow derived macrophages (BMDM) to carry out efferocytosis^8^. We therefore assessed the effects of Nape-pld modulation on efferocytosis by BMDM. Treatment of BMDM with 10 μM of NAPE-PLD inhibitor bithionol (Bith) markedly reduced efferocytosis compared to vehicle treated BMDM, while treatment with 10 μM of either NAPE-PLD activators **VU534** or **VU533** significantly enhanced efferocytosis. (**Figure 7A**). In contrast, treatment with 10 μM compound **VU233** modestly reduced efferocytosis. Given the modest inhibitory effects of compound **VU534** on sEH, we examined the effect of two bona fide sEH inhibitors, AUDA^24^ and TPPU^25^, on efferocytosis. Neither AUDA nor TPPU significantly enhanced efferocytosis (**Figure 7B**), indicating that sEH inhibition was not responsible for the effect of **VU534** on efferocytosis. To further assess the contribution of Nape-pld modulation on efferocytosis, we isolated BMDM from wild-type (WT) and *Napepld*^*-/*-^ (KO) mice and measured the extent of efferocytosis in the presence and absence of activator **VU534**. KO BMDM treated with vehicle (Veh) had significantly reduced efferocytosis compared to WT BMDM treated with Veh (**Figure 7C**). In WT BMDM, treatment with activator **VU534** significantly increased efferocytosis, but in KO BMDM, treatment with activator **VU534** had no effect. Thus, NAPE-PLD appears to play a critical role in maximizing the efferocytosis capacity of macrophages, and activator **VU534** enhances efferocytosis in a NAPE-PLD-dependent manner.

**Figure 7.**
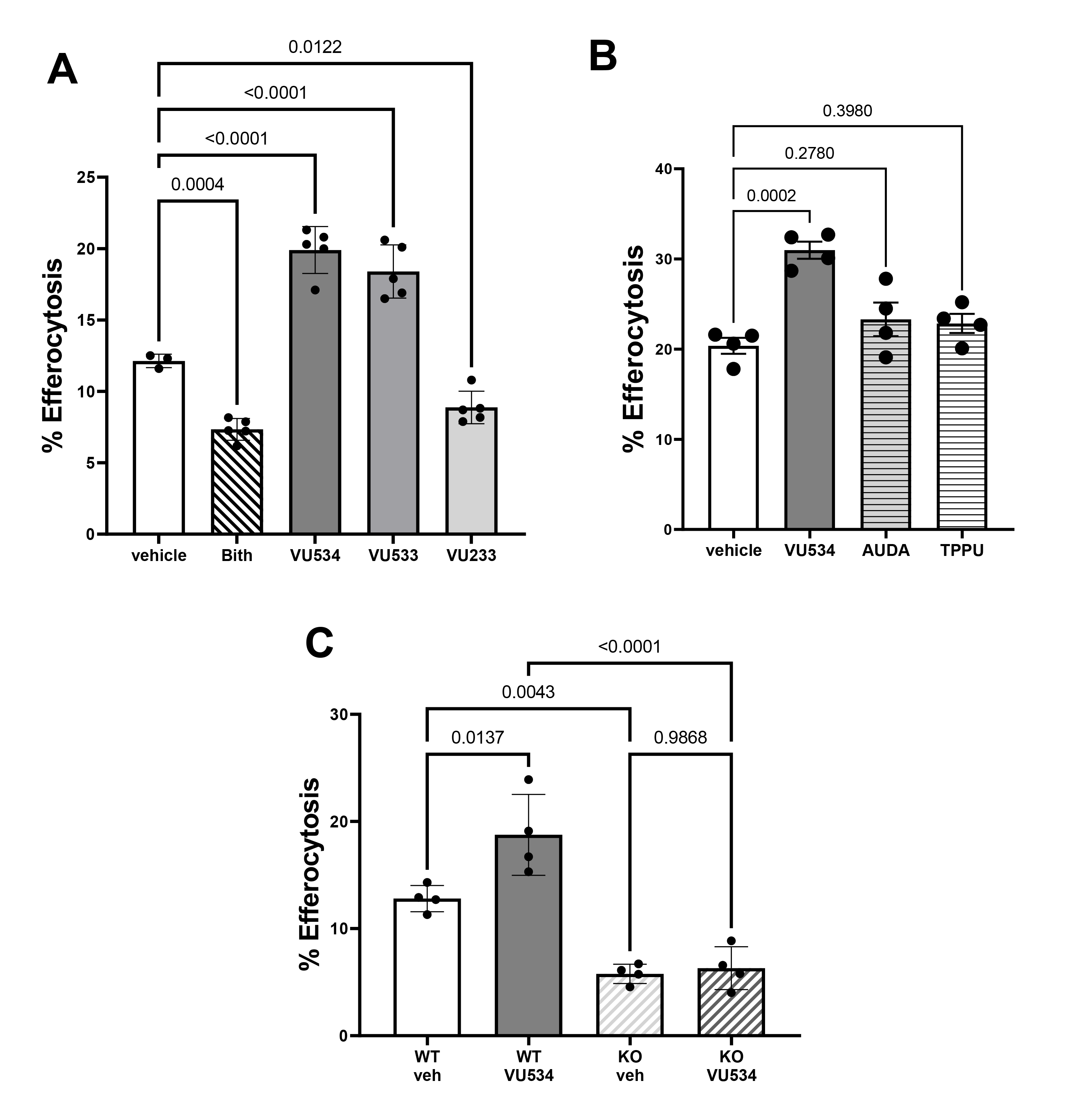
Modulation of NAPE-PLD modulates efferocytosis by macrophages. **A.** BMDM from wild-type mice were treated with 10 μM **VU534, VU533, VU233** or bithionol (Bith) for 6 h prior to initiation of efferocytosis assay. 1-way ANOVA p = 0.0004. Dunnett’s multiple comparison’s test p value shown for each comparison. Data shown from one representative experimental day. All compounds were tested in two to six separate experimental days. **B**. BMDM from wild-type mice were treated with 10 μM **VU534** or with sEH inhibitor AUDA (10 μM) or TPPU (10 μM) for 6 h prior to initiation of efferocytosis assay. 1-way ANOVA p = 0.0004. Dunnett’s multiple comparisons test p value shown for each comparison. Data shown from one representative experimental day of two total experimental days. **C**. BMDM from wild-type (WT) or *Napepld*^-/-^ (KO) mice were treated with vehicle (veh) or 10 μM **VU534** for 6 h prior to initiation of the efferocytosis assay. 1-way ANOVA p<0.0001, Dunnett’s multiple comparison p value shown for each comparison. Data shown from one representative experimental day of three total experimental days.

## Discussion

Our studies demonstrate that select benzothiazole phenylsulfonyl-piperidine carboxamides significantly increase the activity of both mouse and human NAPE-PLD and that increasing NAPE-PLD activity increases efferocytosis by macrophages. The two most potent of the compound series tested, **VU534** and **VU533**, incorporate a para-fluoro-group in the phenylsulfonyl moiety and di-methyl substitution of the benzothiazole aromatic ring. This preliminary SAR suggests that improved activators may be identified from a focused lead optimization campaign. Most of the tested analogs have minimal cytotoxicity against RAW264.7 and HepG2 cells, and their efficacy to enhance cellular Nape-pld activity correlates with their efficacy to enhance activity in the biochemical assay with purified recombinant Nape-pld. Although our lead series share structural features with some previously developed dual FAAH and sEH inhibitors^19^, they show little inhibition of FAAH and only modest inhibition of sEH. Therefore, activators **VU534** and **VU533** should be useful tool compounds to assess the contribution of NAPE-PLD to various biological processes in cultured cells.

A potential role for NAPE-PLD in regulating efferocytosis was suggested by studies showing that PEA, one of the major products of NAPE-PLD, could increase the capacity of macrophages to carry out efferocytosis ^8^. Furthermore, *NAPEPLD* expression was significantly reduced in unstable atherosclerotic plaques from human coronary arteries and impaired efferocytosis has been implicated in the development of these plaques ^8^. Here we show that BMDM derived from *Nape-pld*^*-/-*^ mice have a markedly diminished ability to carry out efferocytosis. Furthermore, we found that inhibiting Nape-pld activity using bithionol phenocopied the effect of *Napepld* deletion. Importantly, treatment of wild-type BMDM with either **VU534** or **VU533** to increase Nape-pld activity markedly enhanced the capacity of BMDM to carry out efferocytosis. This increase in efferocytosis required Nape-pld, as **VU534** failed to enhance efferocytosis by *Napepld*^*-/-*^ BMDM. Together, these results demonstrate the importance of NAPE-PLD in regulating efferocytosis.

Previously described effects of NAEs suggest some mechanisms by which increased NAPE-PLD activity could enhance efferocytosis. PEA acts via Gpr55 to increase the expression of *MerTK*^8^, a receptor that helps macrophages recognize and bind to apoptotic cells^1-2^. Deletion of *MerTK* in macrophages markedly enhances necrotic core expansion in *Apoe*^*-/-*^ mice^26^. OEA acts via PPARα to increase the expression of CD206 and TGFβ, two classic markers of the M2 macrophage phenotype with enhanced efferocytosis^10^. A more complete elucidation of how NAPE-PLD regulates efferocytosis will require a variety of approaches including the use of both NAPE-PLD inhibitors and activators. Efferocytosis is a complex process involving macrophage recognition of so-called “find me” and “eat me” signals, and requires the binding, internalization, and controlled degradation of apoptotic cells, followed by export of their constituent components like cholesterol^1-2, 27^, so the effect of NAPE-PLD modulation on each of these steps needs to be examined. NAPEs exert membrane-stabilizing effects^28-29^ and facilitates the lateral diffusion of cholesterol^30^, while phosphatidic acids exert membrane-bending effects^31^. Therefore, the effect of increased NAPE-PLD activity on membrane topology as a mechanism to enhance efferocytosis also needs to be examined.

Although defective efferocytosis has been implicated in the progression to unstable atherosclerotic plaques^1-2^, whether the enhanced efferocytosis induced by NAPE-PLD activators can translate to improved efferocytosis under atherogenic conditions requires future studies. Previous trials administering OEA, PEA, or NAE-boosting bacteria decrease the size of necrotic cores with atherosclerotic lesions^8, 10-11^. The poor pharmacokinetic properties of OEA and PEA have greatly hampered their clinical use, while engineered bacteria that produce these bioactive lipids in situ still face significant regulatory hurdles for use for humans. While still at a very early stage, our work demonstrates that BT-PSP-based NAPE-PLD activators represent a potential alternative strategy to raise NAE levels and thereby achieve these same effects.

Impaired efferocytosis has been implicated in a number of diseases besides atherosclerosis, including systemic lupus erythematosus, neurodegenerative diseases, retinal degeneration, pulmonary disorders, liver diseases, diabetes, inflammatory bowel disease, colon carcinoma, impaired wound healing, and rheumatoid arthritis^2^. Our results showing a critical role of Nape-pld in regulating efferocytosis suggests the need to examine if Nape-pld expression and activity is reduced in these conditions and whether NAPE-PLD activators can protect against their development or progression. The value of NAPE-PLD activators as a therapeutic intervention may also extend beyond conditions with defective efferocytosis. For instance, Nape-pld expression and NAE levels are rapidly reduced by feeding a high-fat diet^6-8^ and administering OEA or PEA or their precursor NAPEs can markedly blunt the obesity, glucose intolerance, inflammation, and hepatosteatosis that results from these high-fat diets^8-12^. Thus, future testing of NAPE-PLD activators should examine their potential to treat these conditions as well.

## Methods

### Materials

Initial stocks of potential NAPE-PLD modulator compounds were purchased from Life Chemicals and provided by the Vanderbilt HTS screening facility. Additional compounds were synthesized by the Vanderbilt Chemical Synthesis core (**Supplemental Information-Synthesis of Compounds)**. LEI-401, [^2^H_4_]PEA and [^2^H_4_]OEA were purchased from Cayman Chemicals. *N*-palmitoyl-PE, 1,2-dioleoyl-PE and 1,2-dihexanoyl-PE were purchased from Avanti Polar Lipids. PED-A1 was purchased from Invitrogen. Flame-NAPE was synthesized as previously described^18^. [^2^H_4_]*N*-palmitoyl-PE was synthesized using [^2^H_4_] palmitic acid (Cambridge Isotope Laboratories) and 1,2-dioleoyl-PE and *N*-oleoyl-PE was synthesized using 1,2-dihexanoyl-PE and oleoyl chloride (MilliporeSigma) (**Supplemental Information-Synthesis of Compounds)**. Recombinant mouse Nape-pld with a C-terminal hexahistidine tag was expressed in *E. coli* and purified using cobalt affinity beads as previously described^14^. The expression plasmid including the full-length human NAPEPLD gene with a C-terminal hexahistidine tag inserted in a pET plasmid was purchased from VectorBuilder, and the protein expressed and purified in an identical manner to recombinant mouse Nape-pld. The sEH inhibitors TPPU (*N*-[1-(1-Oxopropyl)-4-piperidinyl]-*N*’-[4-(trifluoromethoxy)phenyl]urea) and AUDA (12-[[(tricyclo[3.3.1.13,7]dec-1-ylamino)carbonyl]amino]-dodecanoic acid) were purchased from Cayman Chemical and Sigma Chemicals, respectively.

### Biochemical Nape-pld assays with recombinant enzyme

In vitro fluorescence Nape-pld activity assays using recombinant Nape-pld and either PED-A1 or flame-NAPE as fluorogenic substrate were performed as previously described^14, 18^, except with small modifications as noted below.

For the HTS assays, test compounds were incubated with recombinant enzyme for 1 h prior to adding PED-A1 (final 0.4 μM) mixed with *N*-palmitoyl-dioleoyl-PE (final 3.6 μM) to adjust for the high sensitivity of the Panoptic instrument (WaveFront Biosciences). Assays used black-wall, clear-bottom, non-sterile, and non-treated 384-well plates (Greiner Bio-One 781906). The assay was read in kinetic fluorescence mode on the Panoptic instrument for 4 min and the slope of the signal from 30-100 s was used for analysis. A total of 39,328 compounds from the Vanderbilt Discovery Collection were tested, each at 10 μM. The tested compounds were chosen to represent a structurally diverse selection from the full library. Each 384-well plate included 320 test compound wells and 64 control wells. Unlike our previous pilot screening assay^14^, we used bithionol (10 μM final) in place of lithocholic acid (100 μM final) as the inhibitor control. Before performing high-throughput screening, a checkerboard assay was performed to validate the assay parameters^14, 32^. This yielded a Z’ score of 0.676. Z’ scores were also calculated for each plate during screening, and plates with scores <0.5 were re-run. The average Z’ across all screening plates was 0.52, and the total hit rate was 3.6%. B-scores were calculated from the initial slopes across each plate using WaveGuide software (WaveFront Biosiences).^32^ Modulator hits were defined as compounds with absolute B-scores of 3 or higher. The number of compounds in various B-score ranges were as follows: -21 to -10, 12 compounds; -10 to -5, 221 compounds; -5 to -3, 770 compounds; -3 to 3, 37924 compounds; 3 to 4, 314 compounds; 4 to 6, 70 compounds; 6 to 10, 15 compounds. Activator hits with B-scores of ≥3 were selected for the replication assay and a selection of analogs of the activators.

To identify false hits that modulated fluorescence of the BODIPY moiety indirectly, potential hit compounds were incubated with BODIPY-FL C_5_, a BODIPY-labeled free fatty acid, and measured the effect on fluorescence. One compound directly modulated fluorescence in the absence of enzyme and was therefore eliminated from further evaluation (**Supplemental Figure 7**).

Concentration response curve (CRC) experiments used the same assay conditions as the HTS assay, except that graded concentrations of each test compound were used, with total amount of vehicle (DMSO) kept constant. CRC experiments with purified recombinant mouse Nape-pld were performed on two separate days, with values from each day normalized to vehicle only controls on same plate, and then all normalized values from both days averaged together. CRC experiments with human NAPE-PLD represent value from only a single day, due to limited amounts of this recombinant enzyme.

A similar in vitro fluorescence Nape-pld activity assay was used for LEI-401 and **VU233** competition assays except that this assay was performed using black-walled clear-bottom 96-well plates with 3.5 μM PED-A1 and no *N*-palmitoyl-dioleoyl-PE used as substrate and read in a BioTek Synergy H1 plate reader with the slope of the fluorescence signal from 0-4min (linear phase) used as the assay readout. LEI-401, **VU233**, and **VU534** were incubated for 1 h prior to addition of PED-A1. The Michaelis-Menten study used this same assay except with graded concentration of flame-NAPE.

For LC/MS assays, *N-*oleoyl-PE was added as substrate after 1 h pre-incubation with compound **VU534, VU533, VU233** or vehicle. 90 min after *N-*oleoyl-PE was added, the reaction was quenched by adding 3 volumes of ice-cold methanol containing [^2^H_4_]OEA and [^2^H_4_]*N*-palmitoyl-PE and then 6 volumes of ice-cold chloroform.^33^ The lower phase was dried under nitrogen gas and dissolved in 100 μL mobile phase A. High performance liquid chromatography was performed using a 2.1mm C18 guard column (Phenomenex AJ0-8782), and a rapid gradient ramp. Mobile phase A was 5:1:4 (v/v/v) isopropanol: methanol: water, with 0.2% v/v formic acid, 0.66 mM ammonium formate and 3 μM phosphoric acid included as additives. Mobile phase B was 0.2% (v/v) formic acid in isopropanol. Initial column conditions were 5% mobile phase B, followed by gradient ramp to 95% B over 0.5 min, held at 95% B for 2 min, the returned to initial conditions (5% B) over 1 min. Flow rate throughout was 100 μL/min. Injection volume was 2 μL. The sample injector needle was washed before each injection using a strong wash of methanol, and a weak wash of 1:1:1:1 (v/v/v/v) isopropanol: methanol: acetonitrile: water, with 0.2% formic acid, 0.3 mM ammonium formate, and 0.37 mM phosphoric acid included as additives. Multiple reaction monitoring for the following ions were monitored: OEA [M+H]^+^ : m/z 326.3→ m/z 62.1; [^2^H_4_]OEA [M+H]^+^ : m/z 330.3 → m/z 66.1; *N-*oleoyl-PE [M+NH_4_]^+^ : m/z 693.5 → m/z 308.3 (quantifier), m/z 693.5 → m/z 271.2 (qualifier); [^2^H_4_]*N*-palmitoyl-PE [M+NH_4_]^+^: m/z 1003.8 → m/z 286.3 (quantifier), m/z 1003.8 → m/z 603.5 (qualifier). The ratio of peak height for OEA to [^2^H_4_]OEA was used to calculate to amount of OEA generated and the ratio of peak height for *N-*oleoyl-PE to [^2^H_4_]*N*-palmitoyl-PE was used to calculate the amount of *N-*oleoyl-PE remaining. These values were then used to calculate the OEA / *N*-oleoyl-PE ratio.

### Other biochemical assays

The fluorescence interference assay was performed using the same method as the HTS assay, but with BODIPY-FL C_5_ (ThermoFisher Scientific) used in place of PED-A1. sEH and FAAH activity assays were performed according to the manufacturer’s specifications (Cayman Chemicals).

### Cell-based Nape-pld assays

NAPE-PLD activity was measured in cells as previously described^18^, except that FluoroBrite DMEM (Gibco A1896701) was used as media. For RAW264.7 assays, PED-A1 (3.6 μM final) was used as the substrate with 10 μM orlistat added to inhibit PLA_1_ activity ^18^, while for HepG2 cells, flame-NAPE was used (with no orlistat added).

Cytotoxicity was measured using MTT as previously described^14^ except that the studies used 96-well plates with 100 μL of 0.3 mg/mL MTT solution was added after 24 h of treatment and then replaced after 3 h with 0.1 M HCl in isopropanol. Viability was expressed as percent absorbance at 560 nm relative to vehicle controls.

### Efferocytosis assays

Male C57BL6/j wild-type or Napepld^-/-^ mice^34^ were euthanized with isoflurane and hind legs were removed. Marrow was flushed from femurs and tibias using DMEM containing 4.5 g/L glucose and a 26-gauge needle. Cell suspensions were passed over a 40-µm filter, centrifuged at 500 x g, and resuspended in 50ml of DMEM containing 4.5 g/L glucose, 20% L-cell conditioned media, 10% heat-inactivated FBS, and 1% penicillin/streptomycin. 10 ml of cell suspension was plated into each of five 100-mm dishes and incubated for four days at 37°C and 5% CO_2_. On day four, non-adherent cells and debris were aspirated from the plates and replaced with fresh media. After 7 days of differentiation, cells were harvested for use in experiments.

Assays were performed according to previously established protocols^35 36^. Bone marrow-derived macrophages were seeded at 0.25 × 10^6^ cells/well in a non-tissue culture-treated 24-well plate and allowed to adhere overnight. Macrophages were treated with various compounds at a final concentration of 10 µM or DMSO as a vehicle for 6 hours prior to each experiment. Jurkat cells were exposed to UV light (254nm) for 5 minutes to induce apoptosis and then incubated in a 37°C incubator with 5% CO_2_ for 2 hours. Surveillance staining of these cells routinely yields approximately 80-90% apoptosis (Annexin V^+^) using this method. Apoptotic Jurkat cells were labeled with either CellVue Claret (Millipore Sigma) or Cell Trace Violet (Invitrogen) per the manufacturer’s instructions. After staining, cells were resuspended in macrophage medium at a density of 0.75 × 10^6^ cells/ml and 500 µL of this suspension was added to the drug-containing media on the macrophages to achieve a cell ratio of 3:1 Jurkats:macrophages. After incubating for 45 minutes at 37°C and 5% CO_2_, the medium was aspirated and the macrophages were gently washed twice with PBS to remove unbound apoptotic cells.

Macrophages were then removed from the plate using Cell Stripper (Sigma), washed, resuspended in staining buffer consisting of 2% FBS in PBS with 2mM EDTA, and blocked with anti-mouse CD16/32 antibodies for 15 minutes on ice. After blocking, cells were pelleted and resuspended with F4/80. Cells were incubated for 45 minutes on ice in the dark, then washed and resuspended in staining buffer for analysis. Cells were analyzed using an Attune NxT cytometer (ThermoFisher) and data were analyzed using FlowJo software to quantify the proportion of F4/80^+^ macrophages that co-stained for apoptotic cells (% efferocytosis).

### Statistical Analyses

All statistical analyses and non-linear regression analyses were performed using GraphPad Prism 9 software, except for calculation of B-scores for high throughput screening assays, which were calculated using WaveGuide software (WaveFront Biosiences).

## Acknowledgments

This work was supported by National Institutes of Health Grants P01HL116263 (SSD) and T32GM065086 (JEZ), American Heart Association fellowship 835504 (JEZ), a Discovery Grant from the Vanderbilt Diabetes Center (SSD), the Vanderbilt Institute for Chemical Biology, and the Vanderbilt University Department of Pharmacology Academic Support (SSD). Experiments were performed in the Vanderbilt High-Throughput Screening (HTS) Core Facility with assistance provided by Dr. Paige Vinson, Josh Bauer, and Corbin Whitwell. The Discovery Collection was distributed by the Vanderbilt HTS Core.

The HTS Core receives support from the Vanderbilt Institute of Chemical Biology and the Vanderbilt Ingram Cancer Center (P30 CA68485). The WaveFront Biosciences Panoptic kinetic imaging plate reader is housed and managed within the Vanderbilt HTS Core Facility, an institutionally supported core, and was funded by National Institutes of Health Shared Instrumentation Grant 1S10OD021734. Dr. Ian Romaine assisted in identifying additional compounds in the library with the BT-PSP core structure. Dr. Keri Tallman synthesized and provided the [^2^H_4_]*N*-palmitoyl-PE. The Vanderbilt Mass Spectrometry Research Core receives support from Vanderbilt University.

## Contributions

JEZ performed the majority of experiments including HTS, bioactivity characterization studies, cellular assays, and selectivity studies. He also performed the synthesis of *N*-oleoyl-PE, analyzed data, created figures, interpreted results, and assisted in writing of the initial manuscript. AMAO extracted and cultured BMDM and assisted with efferocytosis studies. CMY performed efferocytosis studies and analysis of data. KK synthesized and characterized BT-PSP analogs and flame-NAPE. ANJ assisted in development and performance of LC/MS assays. MRJ synthesized BT-PSP analogs. ZM assisted with conception of the project, purification of enzyme, LC/MS assays, and cell culture studies. ICS assisted with cellular and LC/MS assays. KM supervised the husbandry of *Napepld*^*-/*-^ and control mice and femur extraction and provided guidance on Nape-pld biology to the project. AGW provided guidance on medicinal chemistry to the project and edited the manuscript. ACD assisted in conception of the project, supervised and performed efferocytosis assays, provided guidance on macrophage biology to the project, created figures, and assisted in writing the initial manuscript. GAS assisted with the conception of the project, supervised the synthesis and characterization of compounds, provided guidance of the project, obtained financial support, and edited the manuscript. SSD conceived and guided the overall project, supervised various studies, obtained financial support, oversaw interpretation of the data, created figures, and wrote the manuscript. All authors reviewed the manuscript.

