## Supplemental Information for "Small Molecule Activation of NAPE-PLD Enhances Efferocytosis by Macrophages"

Jonah E. Zarrow[1], Abdul-Musawwir Alli-Oluwafuyi[1], Cristina M. Youwakim[4], Kwangho Kim[2,3], Andrew N. Jenkins[5], Isabelle C. Suero[1], Margaret R. Jones[2], Zahra Mashhadi[1], Kenneth P. Mackie[6], Alex G. Waterson[1,2,3], Amanda C. Doran[4], Gary A. Sulikowski[1,2,3], and Sean S. Davies[1,3]*.

[1]Department of Pharmacology and [2]Chemistry and the [3]Vanderbilt Institute of Chemical Biology, Vanderbilt University. Nashville, TN. [4]Department of Medicine, Division of Cardiology, Vanderbilt University Medical Center. Nashville, TN, [5]Department of Cell Biology and Physiology, Brigham Young University. Provo, UT. [6]Gill Center and Department of Psychological and Brain Sciences, Indiana University, Bloomington, IN.

### Supplemental Figures

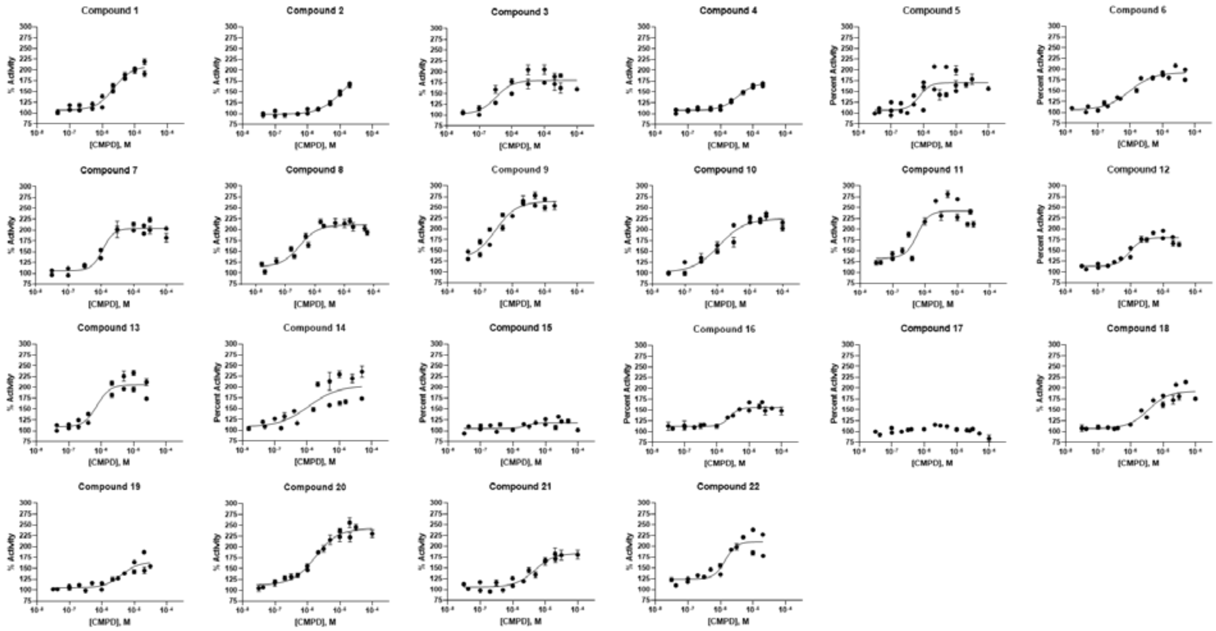


**Supplemental Figure 1. In vitro Nape-pld activity concentration-response curves for series of BT-PSP compounds.** The activity of purified recombinant Nape-pld in the presence of graded concentrations of each entry in the series of BT-PSP compounds in Table 1 was measured using PED-A1 and normalized as % activity, with 100% activity being the average activity without compound (vehicle only). Each compound was tested in triplicate on two or more separate days. Each replicate was normalized to the average of adjacent vehicle only wells, and the three replicates from each day were averaged together. Then these normalized means (± SD) for each experimental day were plotted together on a single graph and the nonlinear curve fit with variable slope (four parameters) generated to determine the EC50 and Emax.

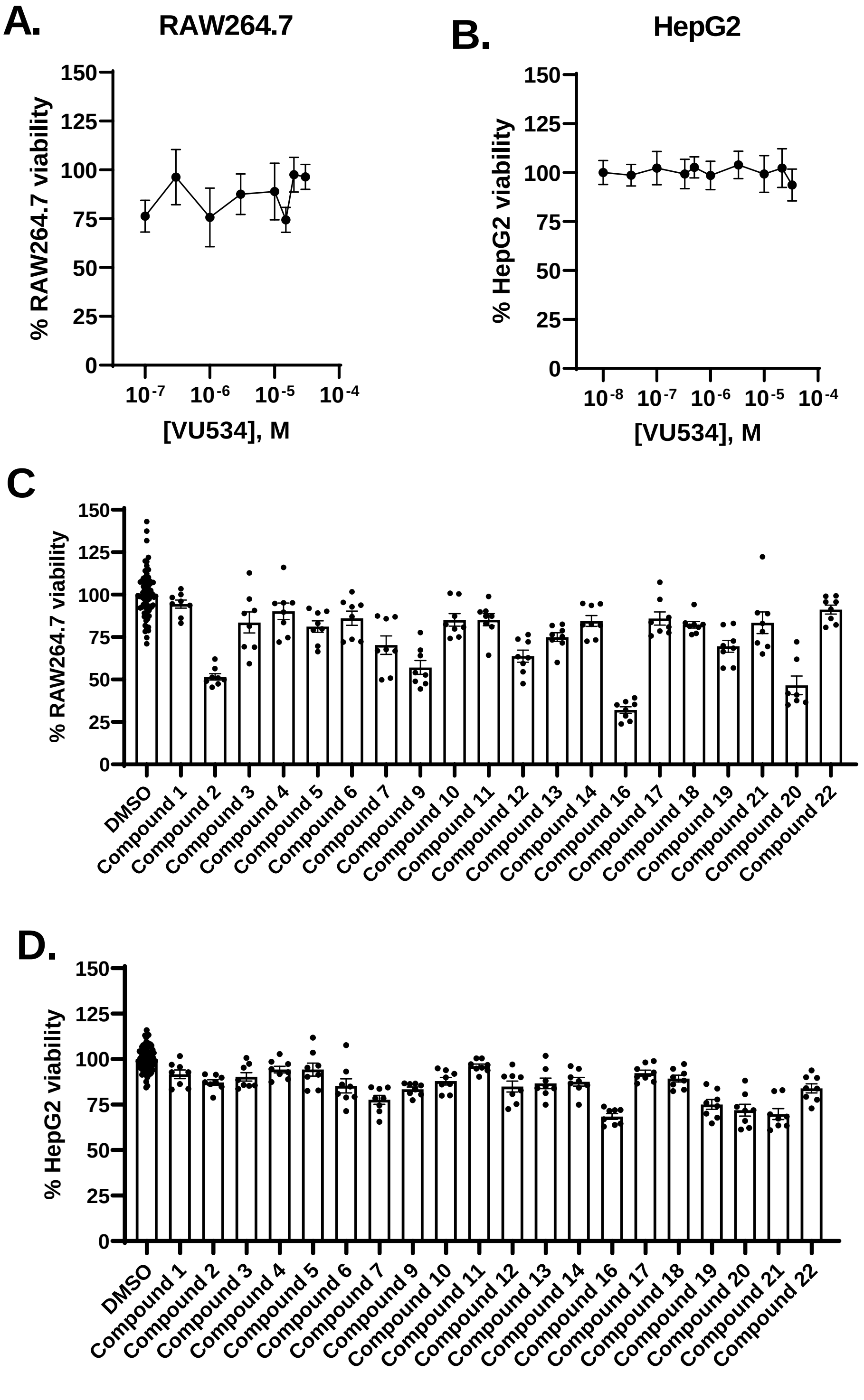


**Supplemental Figure 2. BT-PSPs have minimal cytotoxicity. A.** Effect of graded concentrations of **VU534 (**compound **8)** on cell viability (measured by MTT) of RAW264.7 cells. Values represent mean ± SEM. **B.** Effect of graded concentrations of **VU534 (**compound **8)** on cell viability (measured by MTT) of HepG2 cells. Values represent mean ± SEM. **C.** Effect on

RAW264.7 viability of 30 µM of the series of BT-PSPs (see Table 1 for structures) including compound 9 (**VU533**) and compound 17 (**VU233**). **D.** Effect on HepG2 viability of 30 µM of the same series of BT-PSPs.

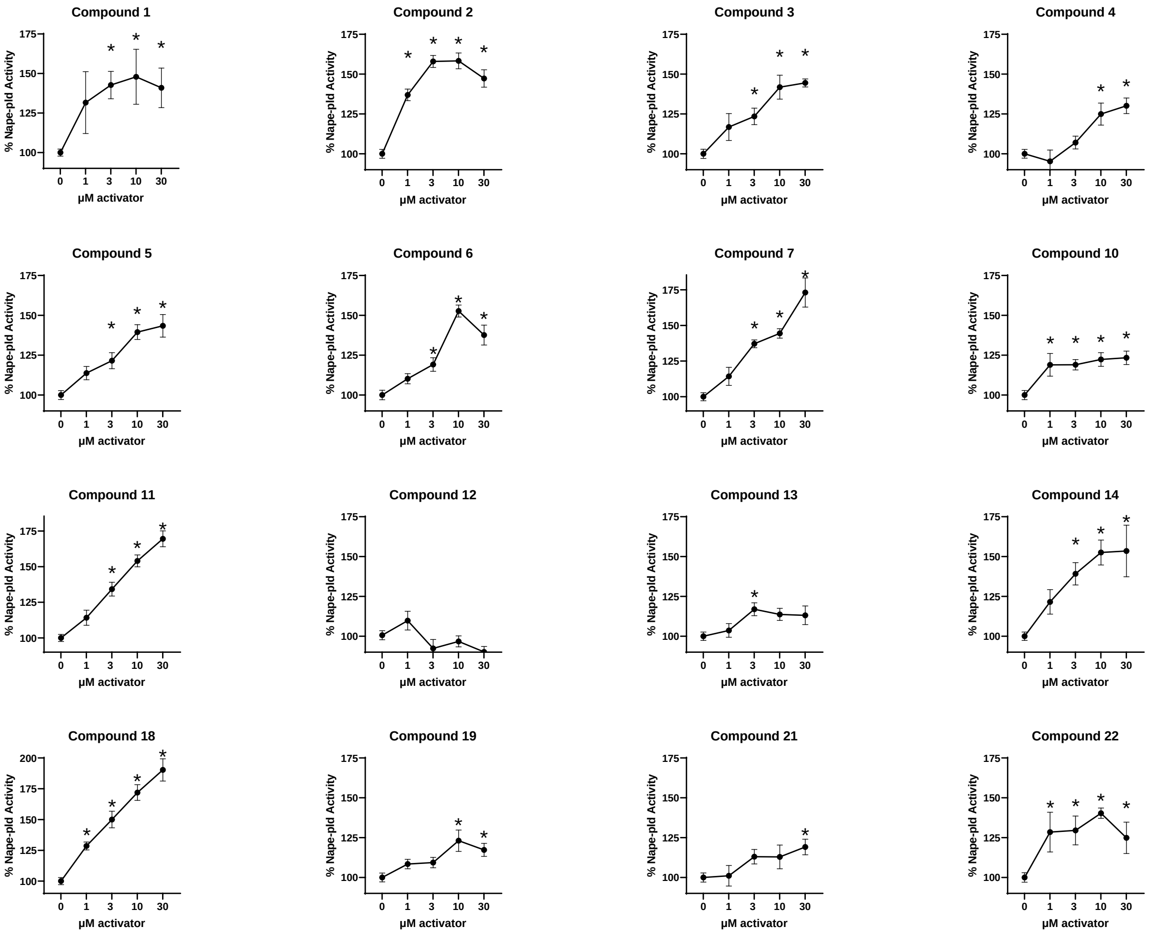


**Supplement Figure 3. BT-PSPs increase the NAPE-PLD activity of RAW264.7 macrophages. A.** Effect of graded concentrations of 19 BT-PSPs (see Table 1 for structures) on Nape-pld activity in RAW264.7 cells, measured using PED-A1. Each compound was tested on at least two separate days and the individual replicates from each day normalized to vehicle control and then combined (mean ± SEM, n = 4-11). 1-way ANOVA $p<0.05$ for all compounds except compounds **12**. * $p<0.05$ vs 0 µM, Dunnet's multiple comparison test.

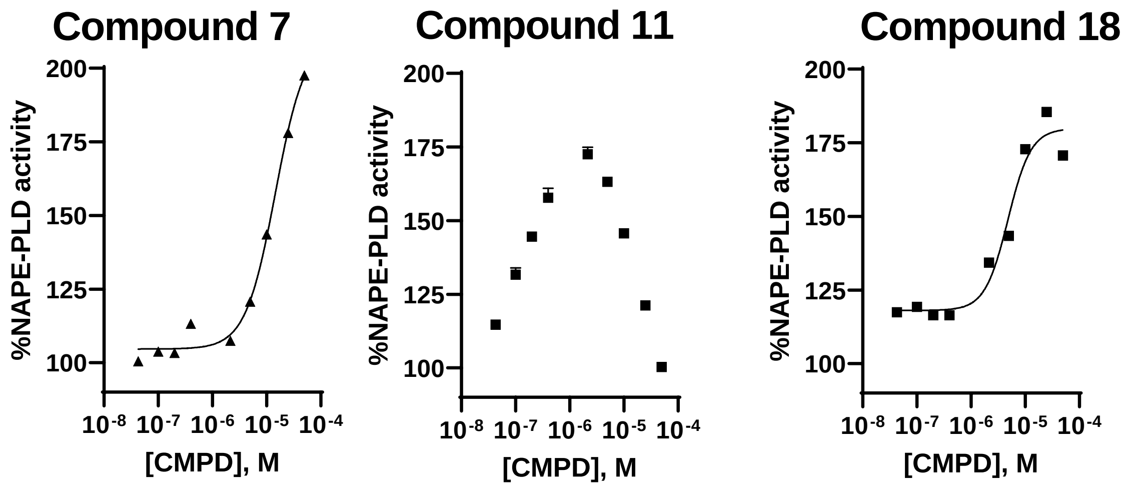


**Supplemental Figure 4. Effects of additional BT-PSPs on activity of recombinant human NAPE-PLD.** Concentration response curves for 3 additional BT-PSPs, compound 7, compound 11, and compound 18 (see Table 1 for structures).

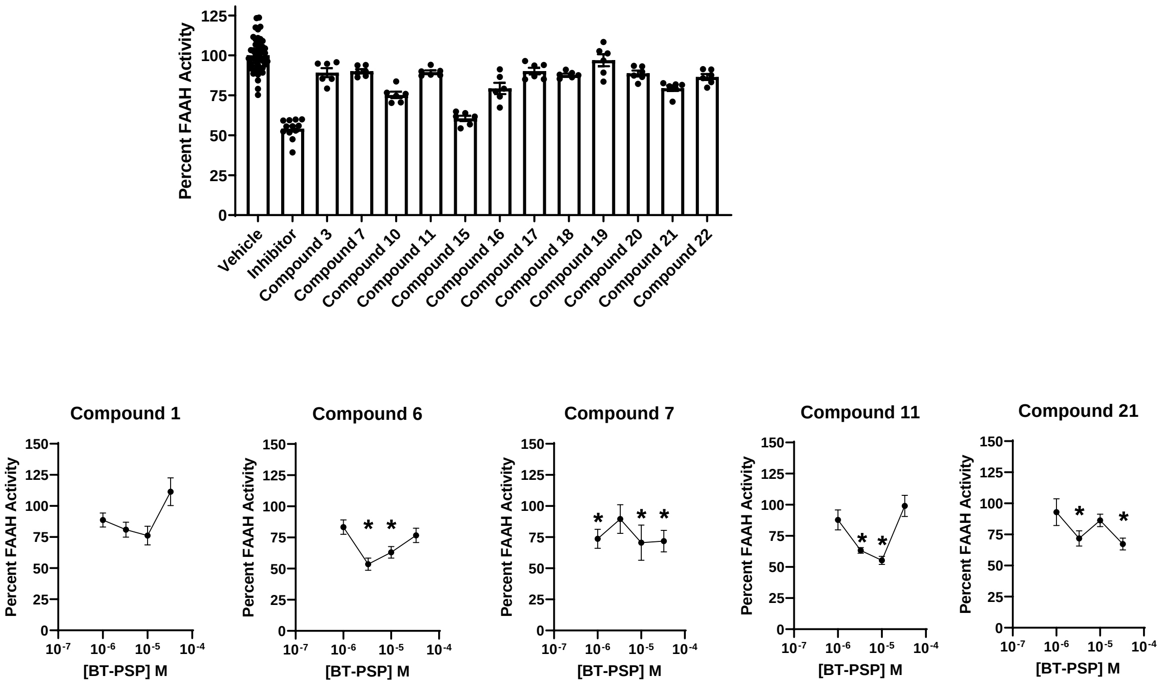


**Supplemental Figure 5.** Effects of various BT-PSP analogs on Fatty Acid Amide Hydrolase activity. Top panel represents initial screen of various BT-PSPs at 30 mM. Lower panel represents concentration response curves for selected compounds.

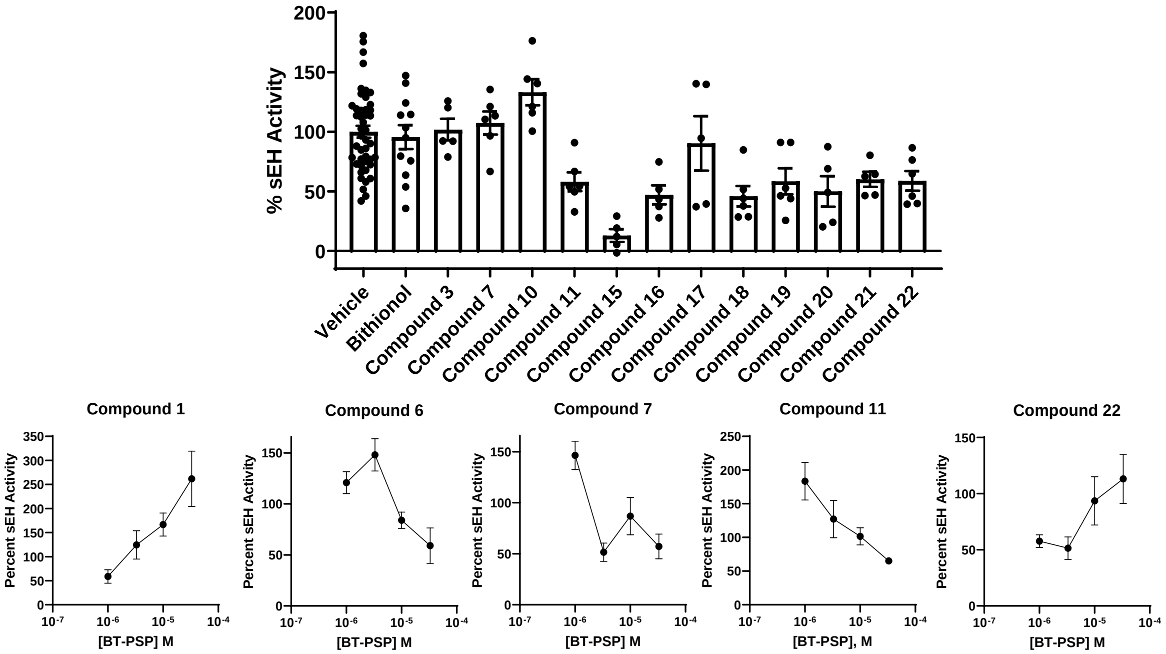


**Supplemental Figure 6.** Effects of various BT-PSPs on soluble epoxide hydrolase (sEH) activity. Top panel represents initial screen of various BT-PSPs at 30 mM. Lower panel represents concentration response curves for selected compounds.

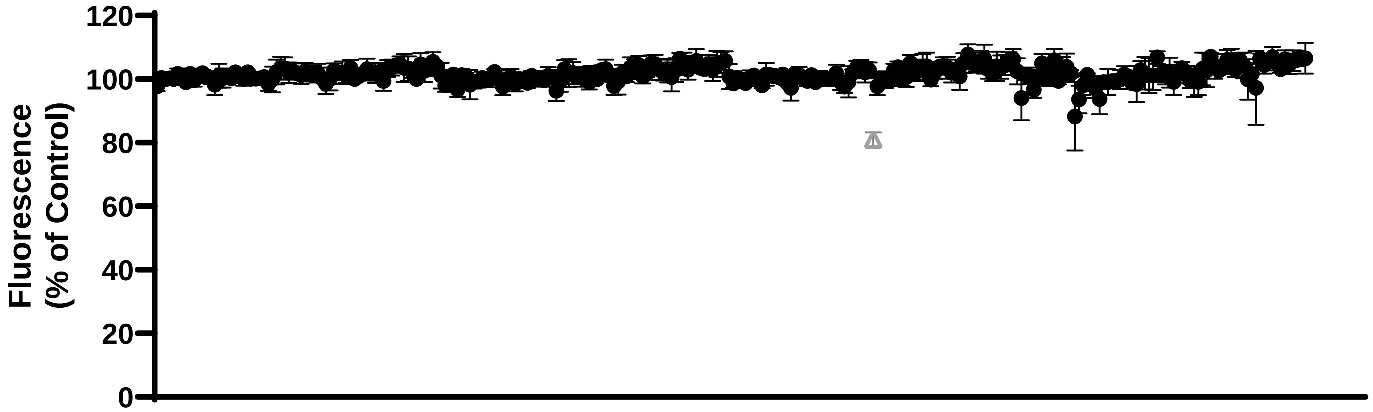


**Supplemental Figure 7. Fluorescence Interference Test.** Effect of candidate modulators on BODIPY fluorescence. Each compound represented by a point. The compound shown as grey triangle significantly differed from control and was eliminated from further evaluation.

**Chemical Synthesis and Characterization**

**Purchased Compounds.** VU484, VU485, VU486, VU488, VU539, VU542, VU517, VU534, VU533, VU575, VU601, VU605 (structures shown in Table 1) were purchased from Life Chemicals. VU542 and VU534 were also synthesized and characterized by the Vanderbilt Chemical Synthesis core as well as the remaining series of additional benzothiazole compounds as described below.

**Synthesis of benzothiazole phenylsulfonyl-piperidine carboxamides**

**General Procedure**: All non-aqueous reactions were performed in flame-dried or oven dried round-bottomed flasks under an atmosphere of argon. Stainless steel syringes or cannula were used to transfer air- and moisture-sensitive liquids. Reaction temperatures were controlled using a thermocouple thermometer and analog hotplate stirrer. Reactions were conducted at room temperature (approximately 23 °C) unless otherwise noted. Flash column chromatography was conducted using silica gel 230-400 mesh. Analytical thin-layer chromatography (TLC) was performed on E. Merck silica gel 60 F254 plates and visualized using UV, and potassium permanganate stain. Yields were reported as isolated, spectroscopically pure compounds.

**Materials**: Solvents were obtained from either an MBraun MB-SPS solvent system or freshly distilled (tetrahydrofuran was distilled from sodium-benzophenone; toluene was distilled from calcium hydride and used immediately; dimethyl sulfoxide was distilled from calcium hydride and stored over 4 Å molecular sieves). Commercial reagents were used as received. The molarity of *n*-butyllithium solutions was determined by titration using diphenylacetic acid as an indicator (average of three determinations).

**Instrumentation**: Semi-preparative reverse phase HPLC was conducted on a Waters HPLC system using a Phenomenex Luna 5 µm C18(2) 100A Axia 250 x 10.00 mm column or preparative reverse phase HPLC (Gilson) using a Phenomenex Luna column (100 Å, 50 x 21.20 mm, 5 µm C18) with UV/Vis detection. Infrared spectra were obtained as thin films on NaCl plates using a Thermo Electron IR100 series instrument and are reported in terms of frequency of absorption ($cm^{-1}$). $^1H$ NMR spectra were recorded on Bruker 400, 500, or 600 MHz spectrometers and are reported relative to deuterated solvent signals. Data for $^1H$ NMR spectra are reported as follows: chemical shift (δ ppm), multiplicity (s = singlet, d = doublet, t = triplet, q = quartet, p = pentet, m = multiplet, br = broad, app = apparent), coupling constants (Hz), and integration. $^{13}C$ NMR spectra were recorded on Bruker 100, 125, or 150 MHz spectrometers and are reported relative to deuterated solvent signals. LC/MS was conducted and recorded on an Agilent Technologies 6130 Quadrupole instrument.

**Compound preparation**

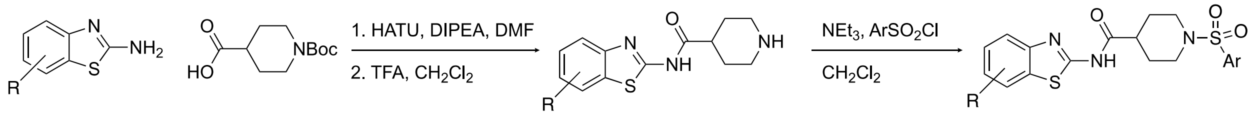


**Supplemental Figure 8.** General preparation of benzothiazole phenylsulfonyl-piperidine carboxamides.

**General amide coupling** To a solution of N-boc-piperidine-4-carboxylic acid (2 g, 13.3 mmol) and HATU (7.6 g, 19.9 mmol) in DMF (40 mL) at 0 °C was added diisopropylethylamine (6.9 mL, 40 mmol) dropwise. After 5 min a solution of a 2-aminobenzothiazole (13.3 mmol) in DMF (5 mL) was added dropwise. The reaction was allowed to warm to ambient temperature and stirred for 24 h. The resulting red-brown solution was quenched with water (20 mL) and extracted with ethyl acetate (3 x 50 mL). The organic extracts were combined and washed with water (20 mL) followed by brine (20 mL). The organic layer was dried ($MgSO_4$), filtered and concentrated in vacuo. The resulting residue was purified by column chromatography.

**General deprotection** To a solution of solution of N-boc-piperidine amide (5.5 mmol) in dichloromethane (20 mL) was added trifluoroacetic acid (1.25 mL, 16.6 mmol). The reaction was heated to reflux and stirred for 16 h. The mixture was cooled to room temperature and concentrated in vacuo. The resulting solid was taken up in a solution of DCM:MeOH (9:1) and neutralized by the addition of 1M NaOH as judged by pH and the mixture extracted with dichloromethane (4 x 50 mL). The organic extracts were combined, washed with brine (10 mL), dried ($MgSO_4$). filtered and concentrated in vacuo. The resulting tan solid was recrystallized from hot chloroform by slow cooling. Further product was achieved by layering of the supernatant with hexanes to give a crystalline solid.

**General sulfonamide formation** A vial (4 mL) equipped with a stir bar was charged with amine (1 equiv) followed by addition of dichloromethane (to a concentration of 100 mM) and triethylamine (1.5 equiv). A solution of arylsulfonyl chloride (1.2 equiv) in dichloromethane was added to the reaction mixture at room temperature. The reaction mixture was maintained for 16 h and diluted with dichloromethane-methanol (9:1). The mixture was washed with saturated $NaHCO_3$ (aq.) followed by water. The organic layer was dried ($MgSO_4$) and concentrated in vacuo. The residue was purified by recrystallization from hot chloroform.

**VU542 (entry 6)** From 4-Chloro-benzenesulfonyl chloride (46 mg, 0.22 mmol) according to the general procedure. 42 mg, 51% of the product was obtained as an off-white crystalline solid: 1H NMR ($d_6$-DMSO, 400 MHz): δ 7.79-7.72 (m, 5H), 7.22 (d, *J* = 7.2 Hz, 1H), 7.28 (t, *J* = 8.8 Hz, 1H), 4.07 (q, *J* = 5.2 Hz, 1H), 3.68-3.65 (m, 2H), 2.54 (s, 3H), 2.35 (dt, d, *J* = 11.6 Hz, 2.4 Hz, 2H), 1.98-1.90 (m, 2H), 1.65 (dq, *J* = 13.2 Hz, 4.0 Hz, 2H); LCMS calc'd for C20H20ClN3O3S2 $[M+H]^+$: 449.9, measured 450.1.

**VU534 (entry 8)** From 4-Fluoro-benzenesulfonyl chloride (97 mg, 0.50 mmol) according to the general procedure. 94 mg, 84% of the product was obtained as an off-white crystalline solid: 1H NMR (CDCl3, 400 MHz): δ 7.70 (dd, *J* = 9.2 Hz, 5.2 Hz, 1H), 7.69 (t, *J* = 4.8 Hz, 1H), 7.28 (bs, 1H), 7.22 (t, *J* = 8.8 Hz, 2H), 6.56 (s, 1H), 3.66 (dd, *J* = 8.4 Hz, 3.2 Hz, 2H), 2.52 (s, 3H), 2.32-2.26 (m, 1H), 2.25 (s, 3H), 1.98 (dt, *J* = 11.6 Hz, 2.4 Hz, 2H), 1.86 (dq, *J* = 11.6 Hz, 4.0 Hz, 2H), 1.74 (dd, *J* = 13.6 Hz, 3.2 Hz, 2H) ; 13C NMR (CDCl3, 400 MHz): δ 172.6, 166.4, 163.9, 159.5, 147.2, 136.4, 132.3, 132.2, 131.7, 130.2, 130.1, 128.8, 126.2, 117.6, 116.4, 116.2, 44.9, 41.6, 27.6, 21.3, 20.5 ; LCMS calc'd for C21H22FN3O3S2 $[M+H]^+$: 447.5, measured 448.1.

**VU231 (entry 14)** From 4-Fluoro-benzenesulfonyl chloride (97 mg, 0.50 mmol) according to the general procedure. 65 mg, 60% of the product was obtained as an off-white crystalline solid: 1H NMR (CD3OD, 400 MHz): δ 7.88 (dd, *J* = 8.8 Hz, 4.8 Hz, 1H), 7.87 (t, *J* = 4.8 Hz, 1H), 7.65 (d, *J* = 7.6 Hz, 1H), 7.37 (dt, *J* = 8.8 Hz, 2.0 Hz, 2H), 7.22 (d, *J* = 6.4 Hz, 1H), 7.19 (q, *J* = 7.6 Hz, 1H), 3.82 (dd, *J* = 8.4 Hz,

3.2 Hz, 2H), 2.63 (s, 3H), 2.6-2.45 (m, 3H), 2.01 (dd, *J* = 13.6 Hz, 3.2 Hz, 2H), 1.87 (dq, *J* = 11.6 Hz, 4.0 Hz, 2H); LCMS calc'd for C20H20FN3O3S2 $[M+H]^+$: 433.5, measured 434.1.

**VU205 (entry 15)** From 4-fluoro-2-methylbenzenesulfonyl chloride (48 mg, 0.23 mmol) according to the general procedure. 53 mg, 64% of the product was obtained as a beige crystalline solid: 1H NMR ($d_6$-DMSO, 400 MHz): δ 7.95 (d, *J* = 8.1 Hz, 1H), 7.88 (dd, *J* = 8.8 Hz, 5.9 Hz, 1H), 7.71 (d, *J* = 8.0 Hz, 1H), 7.42 (t, *J* = 7.8 Hz, 1H), 7.37 (dd, *J* = 9.9 Hz, 2.7 Hz, 1H), 7.28 (m, 2H), 3.63 (d, *J* = 12.5 Hz, 2H), 2.68 (m, 3H), 2.57 (s, 3H), 1.94 (d, *J* = 13.6 Hz, 2H), 1.62 (qd, *J* = 11.9 Hz, 3.6 Hz, 2H); 13C NMR ($d_6$-DMSO, 400 MHz): δ 173.9, 165.7, 163.2, 158.2, 148.9, 141.5, 133.1, 132.5, 131.8, 126.5, 123.9,122.1, 120.9, 120.1, 119.9, 113.7, 75.1, 44.7, 41.0, 27.86, 20.6; LCMS calc'd for C20H20FN3O3S2 $[M+H]^+$: 433.5, measured 434.1.

**VU212 (entry 16)** From 4-fluoro-2-methylbenzenesulfonyl chloride (48 mg, 0.23 mmol) according to the general procedure. 44 mg, 54% of the product was obtained as an off-white crystalline solid: $^1$H NMR ($d_6$-DMSO, 400 MHz): δ 7.88 (dd, *J* = 8.9 Hz, 5.9 Hz, 1H), 7.76 (d, *J* = 7.7 Hz, 1H), 7.36 (dd, *J* = 9.6 Hz, 2.5 Hz, 1H), 7.26 (td, *J* = 8.4 Hz, 2.9 Hz, 1H), 7.23 (d, *J* = 7.1 Hz, 1H), 7.17 (t, *J* = 7.6 Hz, 1H), 3.63 (d, *J* = 12.8 Hz, 2H), 2.66 (m, 3H), 2.57 (s, 3H), 2.54 (s, 3H), 1.93 (d, *J* = 12.8 Hz, 2H), 1.62 (qd, *J* = 12.6 Hz, 3.4 Hz, 2H); $^{13}$C NMR ($d_6$-DMSO, 400 MHz): δ 173.9, 157.4, 147.9, 141.5, 133.1, 132.5, 131.5, 130.1, 126.9, 123.8, 120.0, 119.4, 113.7, 74.9, 44.7, 41.0, 27.9, 20.6, 18.4; LCMS calc'd for C21H22FN3O3S2 $[M+H]^+$: 447.5, measured 448.1.

**VU233 (entry 17)** From 3-chloro-4-fluorobenzenesulfonyl chloride (53 mg, 0.23 mmol) according to the general procedure. 59 mg, 68% of the product was obtained as a white crystalline solid: 1H NMR ($d_6$-DMSO, 400 MHz): δ 7.98 (dd, *J* = 6.8 Hz, 2.2 Hz, 1H), 7.94 (d, *J* = 7.9 Hz, 1H), 7.80 (ddd, *J* = 8.7 Hz, 2.4 Hz, 2.2 Hz, 1H), 7.71 (m, 2H), 7.41 (t, *J* = 7.8 Hz, 1H), 7.28 (t, *J* = 7.8 Hz, 1H), 3.68 (d, *J* = 11.9 Hz, 2H), 2.54 (tt, *J* = 11.5 Hz, 3.8 Hz, 1H), 2.41 (td, *J* = 11.8 Hz, 2.1 Hz, 2H), 1.95 (dd, *J* = 13.4 Hz, 2.4 Hz, 2H), 1.65 (qd, *J* = 12.3 Hz, 3.9 Hz, 2H); $^{13}$C NMR ($d_6$-DMSO, 400 MHz): δ 173.9, 159.0, 158.3, 148.9, 133.6, 131.8, 130.3, 129.3, 126.5, 123.9, 122.1, 121.6, 121.4, 120.8, 118.7, 118.5, 75.2, 45.7, 40.6, 27.6; LCMS calc'd for C19H17ClFN3O3S2 $[M+H]^+$: 453.9, measured 454.1.

**VU209 (entry 18)** From 3-chloro-4-fluorobenzenesulfonyl chloride (53 mg, 0.23 mmol) according to the general procedure. 32 mg, 38% of the product was obtained as a white crystalline solid: $^1$H NMR ($d_6$-DMSO, 400 MHz): δ 7.99 (dd, *J* = 6.8 Hz, 2.2 Hz, 1H), 7.81 (m, 1H), 7.75 (d, *J* = 7.7 Hz, 1H), 7.71 (t, *J* = 8.9 Hz, 1H), 7.23 (d, *J* = 6.9 Hz, 1H), 7.17 (t, *J* = 7.8 Hz, 1H), 3.69 (d, *J* = 11.9 Hz, 2H), 2.55 (s, 3H), 2.54 (m, 1H), 2.40 (t, *J* = 12.0 Hz, 2H), 1.95 (d, *J* = 12.6 Hz, 2H), 1.64 (qd, *J* = 12.6 Hz, 3.4 Hz, 2H); $^{13}$C NMR ($d_6$-DMSO, 400 MHz): LCMS calc'd for C20H19ClFN3O3S2 $[M+H]^+$: 467.9, measured 468.1.

**VU227 (entry 19)** From (1-methyl-1H-pyrazol-4-yl)sulfonyl chloride (41 mg, 0.23 mmol) according to the general procedure. 32 mg, 42% of the product was obtained as a white crystalline solid: 1H NMR ($d_6$-DMSO, 400 MHz): δ 8.34 (s, 1H), 7.94 (d, *J* = 7.8 Hz, 1H), 7.79 (s, 1H), 7.70 (d, *J* = 7.7 Hz, 1H), 7.40 (t, *J* = 7.7 Hz, 1H), 7.27 (t, *J* = 7.7 Hz, 1H), 3.91 (s, 3H), 3.57 (d, *J* = 11.6 Hz, 2H), 2.52 (m, 1H), 2.27 (td, *J* = 11.8 Hz, 2.1 Hz, 2H), 1.97 (dd, *J* = 13.4 Hz, 2.4 Hz, 2H), 1.69 (qd, *J* = 12.3 Hz, 3.9 Hz, 2H); LCMS calc'd for C17H19N5O3S2 $[M+H]^+$: 405.5, measured 406.1.

**VU210 (entry 20)** From (1-methyl-1H- pyrazol-4-yl)sulfonyl chloride (41 mg, 0.23 mmol) according to the general procedure. 43 mg, 56% of the product was obtained as a white crystalline solid: 1H NMR ($d_6$-DMSO, 400 MHz): δ 8.34 (s, 1H), 7.80 (s, 1H), 7.75 (d, *J* = 7.6 Hz, 1H), 7.23 (d, *J* = 7.2 Hz, 1H), 7.17 (t, *J* = 7.6 Hz, 1H), 3.91 (s, 3H), 3.56 (d, *J* = 11.9 Hz, 2H), 2.54 (s, 3H), 2.51 (m, 1H), 2.25 (t, *J* = 11.8 Hz, 2H), 1.96 (d, *J* = 12.6 Hz, 2H), 1.68 (qd, *J* = 12.6 Hz, 3.4 Hz, 2H); $^{13}$C NMR ($d_6$-DMSO, 400 MHz): δ 174.0, 138.9, 133.7, 131.5, 130.1, 126.9, 123.8, 119.4, 116.3, 75.0, 45.9, 40.7, 27.5, 18.4; LCMS calc'd for C18H21N5O3S2 $[M+H]^+$: 419.5, measured 420.2.

**VU203 (entry 21)** From pyridine-3-sulfonyl chloride (41 mg, 0.23 mmol) according to the general procedure. 46 mg, 53% of the product was obtained as a white crystalline solid: $^1$H NMR ($d_6$-DMSO, 400 MHz): δ 8.92 (d, *J* = 2.1 Hz, 1H), 8.89 (dd, *J* = 4.9 Hz, 1.5 Hz, 1H), 8.19 (dt, *J* = 8.2 Hz, 2.1 Hz, 1H), 7.93 (d, *J* = 7.8 Hz, 1H), 7.70 (m, 2H), 7.40 (t, *J* = 7.7 Hz, 1H), 7.27 (t, *J* = 7.7 Hz, 1H), 3.70 (d, *J* = 11.9 Hz, 2H), 2.56 (tt, *J* = 11.5 Hz, 3.8 Hz, 1H), 2.42 (td, *J* = 11.8 Hz, 2.1 Hz, 2H), 1.95 (dd, *J* = 13.4 Hz, 2.4 Hz, 2H), 1.63 (qd, *J* = 12.3 Hz, 3.9 Hz, 2H); $^{13}$C NMR ($d_6$-DMSO, 400 MHz): δ 174.0, 158.5, 154.1, 148.9, 148.1, 135.9, 132.7, 131.8, 126.4, 124.9, 123.8, 122.0, 120.8, 75.0, 45.5, 40.6, 27.6; LCMS calc'd for C18H18N4O3S2 $[M+H]^+$: 402.5, measured 403.1.

**VU0934211 (entry 22)** From pyridine-3-sulfonyl chloride (41 mg, 0.23 mmol) according to the general procedure. 26 mg, 34% of the product was obtained as a beige crystalline solid: $^1$H NMR ($d_6$-DMSO, 400 MHz): δ 8.92 (d, *J* = 2.2 Hz, 1H), 8.90 (dd, *J* = 4.9 Hz, 1.4 Hz, 1H), 8.18 (dt, *J* = 8.0 Hz, 1.9 Hz, 1H), 7.73 (d, *J* = 7.7 Hz, 1H), 7.70 (dd, *J* = 7.8 Hz, 4.9 Hz, 1H), 7.22 (d, *J* = 7.1 Hz, 1H), 7.17 (t, *J* = 7.5 Hz, 1H), 3.71 (d, *J* = 12.0 Hz, 2H), 2.55 (m, 1H), 2.54 (s, 3H), 2.42 (t, *J* = 11.8 Hz, 2H), 1.94 (d, *J* = 12.6 Hz, 2H), 1.64 (qd, *J* = 12.6 Hz, 3.4 Hz, 2H); $^{13}$C NMR ($d_6$-DMSO, 400 MHz): δ 173.8, 157.4, 154.1, 148.1, 147.9, 136.0, 132.7, 131.5, 130.1, 126.9, 124.9, 123.8, 119.4, 45.5, 27.6, 18.4; LCMS calc'd for C19H20N4O3S2 $[M+H]^+$: 416.5, measured 417.1.

### NAPE synthesis

**1,2-dihexanoyl-*sn*-glycero-3-phospho-*N*-oleoyl-ethanolamine (*N*-oleoyl-PE)**

2.5 mL of dry chloroform was added to a 25mL round-bottom flask. To that, 1,2-dihexanoyl-*sn*-glycero-3-phosphoethanolamine (164.6 µL, 4 µmol, Avanti Polar Lipids), triethylamine (1.12 µL, 8 µmol), and oleoyl chloride (1.49 µL, 4 µmol, Millipore Sigma) were added. The reaction was stirred for 19 h at room temperature. After the reaction was completed, product was collected using a modified Folch extraction (final extraction solvent composition 6:2:1, chloroform, methanol, saturated $NaHCO_3$ solution v/v/v). The chloroform (lower) layer was transferred to a glass 10 mL sample tube and dried under $N_2$ gas. Then, the dried product was dissolved in 2 mL deionized water and 4mL of ice-cold Folch solution (2:1 chloroform:methanol). This mixture was vortexed for 5 min and incubated on ice for 30 min. The organic and aqueous layers were separated by centrifugation (500xg, 5 min, 4 °C), and the chloroform (lower) layer was saved while the aqueous was discarded. The chloroform layer was dried under gaseous $N_2$ and re-dissolved in 1 mL of chloroform. This was passed over a Sep-pak plus silica gel cartridge (Waters #WAT036580). The column was washed with 8 mL of 1:9 methanol:chloroform, and then the *N*-oleoyl-PE eluted with 8 mL of Folch solution. Eluted *N*-oleoyl-PE was dried under gaseous $N_2$ and re-dissolved in 800 µL of chloroform. Product was stored in an amber glass vial at -20 °C. LC/MS was conducted and recorded on an ThermoFinnigan

Quantum electrospray ionization triple quadrupole mass spectrometer in positive ion mode. LCMS calc'd for C35H67NO9P+ [M+H]+ 676.5, measured 676.6.

**1,2-dioleoyl *sn*-glycero-3-phospho-*N*-$[^{2}H_{4}]$oleoyl-ethanolamine ($[^{2}H_{4}]$*N*-palmitoyl-PE)**

HOBT (8.0 mg, 0.052 mmol) and EDC-HCl (10 mg, 0.052 mmol) were added to a solution of 1,2-dioleoyl-sn-3-glycerophosphoethanolamine (25 mg, 0.034 mmol, Avanti Polar Lipids) and 7,7,8,8-d4-palmitic acid (9.0 mg, 0.035 mmol, Cambridge Isotopes) in CHCl3 (1 mL). After allowing to react overnight, the reaction mixture was diluted with CHCl3/MeOH (30 mL) and washed with saturated NH4Cl (10 mL) and concentrated. The product was purified by column chromatography on silica gel (10% MeOH/CH2Cl2) and isolated as a white sticky solid (31 mg, 94%). 1H NMR (CDCl3)  5.36 -5.27 (m, 4H), 5.22-5.18 (m, 1H), 4.37-4.29 (m, 1H), 4.13-4.09 (m, 2H), 3.92-3.90 (m, 3H), 3.50-3.43 (m, 2H), 2.72 (br s, 1H), 2.32-2.24 (m, 5H), 2.20-2.14 (m, 2H), 1.97 (app q, 8H, J = 5.9 Hz), 1.62-1.50 (m, 6H), 1.27-1.23 (m, 60 H), 0.85 (t, 9H, J = 6.5 Hz).

LC/MS was conducted and recorded on an ThermoFinnigan Quantum electrospray ionization triple quadrupole mass spectrometer in positive ion mode. LCMS calc'd for C57H105D4NO9P+ [M+H]+ 986.8, measured 986.8.
